# Tumor Immune Cell Targeting Chimeras (TICTACs) reprogram tumor-associated macrophages

**DOI:** 10.1101/2023.12.06.570444

**Authors:** Mariko Morimoto, David S. Roberts, Ru M. Wen, Ell Handy, Eric E. Peterson, Grace M. Stepek, Nicholas A. Till, James D. Brooks, Carolyn R. Bertozzi

## Abstract

Immune cells in the tumor microenvironment are not only powerful regulators of immunosuppression and tumorigenesis, but also a dominant cell population, with tumor-associated macrophages (TAMs) comprising up to 50% of solid tumor mass. Immunotherapies such as immune checkpoint inhibitors derive efficacy from this cancer-immune interface; however, immune-related adverse events from systemic blockade remain a major challenge. To address this need for potent, tumor-specific immunotherapies, we developed Tumor-Immune Cell TArgeting Chimeras (TICTACs) that selectively deplete immune checkpoint receptors such as SIRPα from TAM surfaces. These chimeras consist of a synthetic ligand targeting CD206, a TAM marker, conjugated to a non-blocking antibody that binds without inhibiting the checkpoint receptor. By engaging CD206, which constitutively recycles between the plasma membrane and early endosomes, TICTACs drive robust checkpoint degradation in CD206^high^ macrophages, with no effect on CD206^low^ cells. This decoupling of antibody selectivity from blocking function presents a new paradigm for tumor-specific immunotherapies.

## Introduction

Over the past 10 years, cancer immune therapy has significantly improved the therapeutic landscape for multiple cancer subtypes^1^. In particular, the PD-1 blocking immune checkpoint inhibitors (ICIs) pembrolizumab and nivolumab have become a mainstay in the first-line treatment of melanoma and non-small cell lung cancer (NSCLC)^2^. These antibody-based therapeutics act through blocking the inhibitory receptor PD-1 that is present on immune cells, thus potentiating the body’s immune response against the tumor. Nonetheless, significant challenges remain with respect to efficacy in diverse cancer subtypes as well as target-related side effects that can be dose limiting. FDA-approved ICI therapies against PD-1, CTLA-4^3^, and LAG-3^4^, target cell surface markers without further specificity for immune cell subtype, leading to pleiotropic effects including unwanted immune-related adverse events (irAEs)^5–8^. It has been proposed that ICIs with increased immune cell and tissue specificities would provide a path towards decreased irAEs without sacrificing efficacy. With this principle in mind, we sought to develop a platform for tumor-immune cell targeting with the eventual goal of interfacing with existing ICI technologies.

We focused on tumor-associated macrophages (TAMs), which represent a highly abundant immune cell type within the tumor microenvironment (TME)^9–11^. Unlike classically activated M1 macrophages, TAMs predominantly adopt the M2-like, pro-tumor phenotype in response to cytokines such as IL-4 within the TME. These M2-like macrophages notably aid cancer cell metastasis, angiogenesis, and proliferation via diverse anti-inflammatory mechanisms.^12^ CD206, also known as the macrophage mannose receptor (MMR), is one of the most prominent M2 markers, and its expression is negatively correlated with overall survival of patients with solid tumors (**Fig. 1a**)^13–15^.

**Figure 1.**
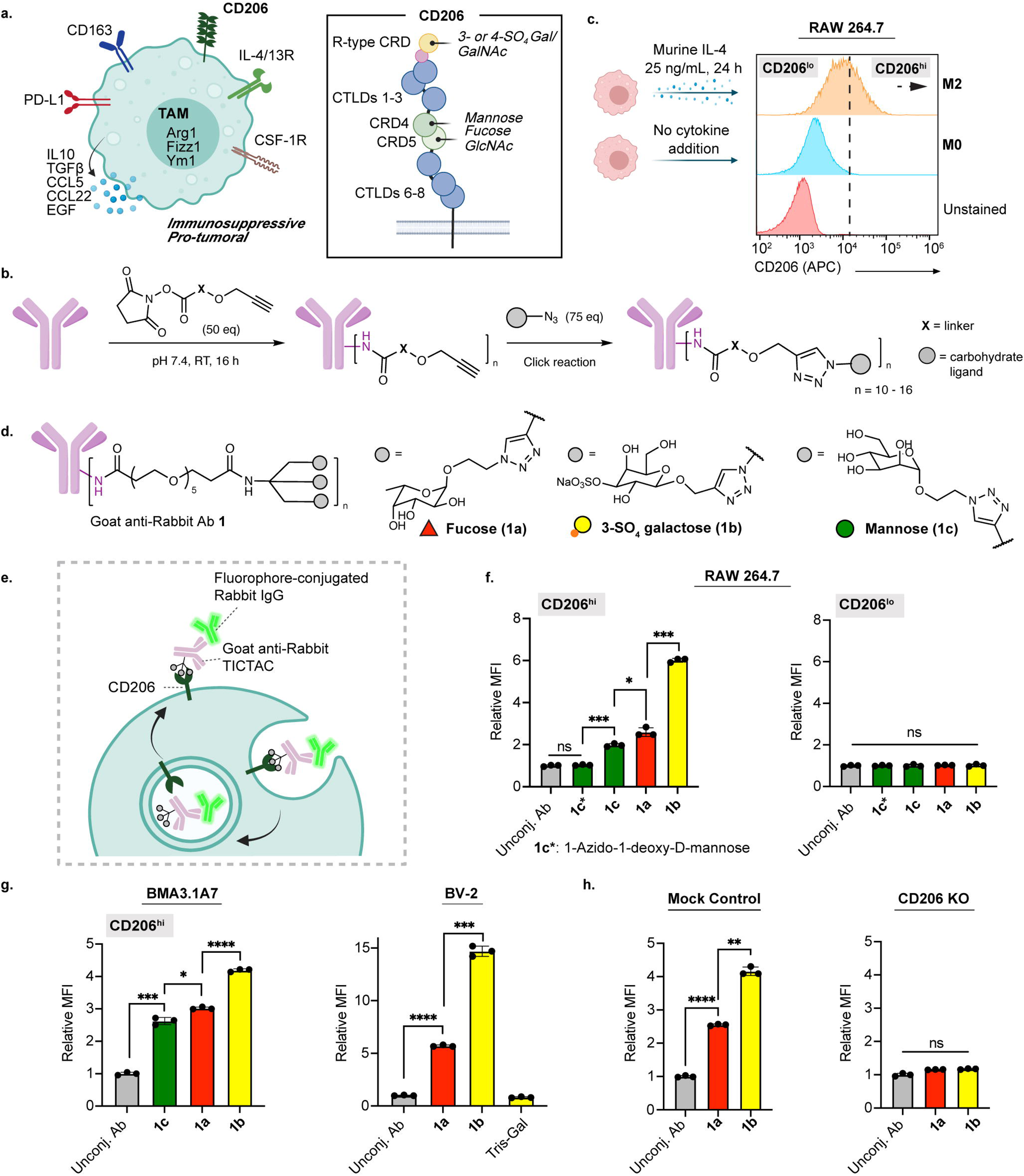
TICTACs target CD206 on M2-polarized macrophages. **a,** CD206 is upregulated on the surface of M2-polarized TAMs and binds multivalent mannose, fucose, and glucosamine via its C-type lectin-like domains (CTLDs) and sulfated galactose and galactosamine via its N-terminal R-type carbohydrate recognition domain (R-type CRD). **b,** Two-step protocol for generating TICTACs, where X represents the linker, and the gray sphere represents the carbohydrate ligand. **c,** Murine macrophages (RAW264.7) can be M2-polarized upon treatment with IL-4, leading to increased expression of CD206. CD206^lo^ and CD206^hi^ populations are shown live cell flow cytometry. **d,** Tris-dendron scaffold enables CD206-targeting with fucose, 3-SO_4_ galactose, and mannose ligands **1a-c**, respectively. **e,** TICTAC-mediated internalization of fluorophore-conjugated rabbit IgG into M2-polarized macrophages. **f,** Fold change in mean fluorescence intensity (MFI) relative to the control for M2-polarized RAW264.7 cells incubated at 37 °C for 3 h with 25 nM rabbit IgG-488 and 25 nM unconjugated goat-anti-rabbit antibody (lacking carbohydrate ligands) or various goat-anti-rabbit TICTACs. MFI was determined by live cell flow cytometry. Data are meanL±Ls.d. of *n*L=L3 biological replicates. **g,** MFI relative to unconjugated antibody control for M2 polarized murine macrophage cell line BMA3.1A7 and microglia cell line BV-2. Data are meanL±Ls.d. of *n*L=L3 biological replicates. **h,** Knocking out CD206 ablates **1a** and **1b**-mediated uptake in RAW264.7 cells. Data are meanL±Ls.d. of *n*L=L3 biological replicates. For **f** and **g**, *P* values were determined by one-way ANOVA followed by Tukey’s post-hoc test for multiple comparisons, and by a two-sided unpaired t-test for **h**. Statistical significance was defined as *P*<0.05, and the asterisks * indicates *P*<0.05, ** indicates *P*<0.01, *** indicates *P*<0.001, and **** indicates *P*<0.0001.

Here we developed tumor immune cell targeting chimeras (TICTACs), which are bifunctional small molecule-antibody conjugates that selectively bind CD206 on macrophages. Through a click-based two-step approach, we created a modular platform that enables facile optimization of the multivalent display of the small molecule ligand and show that these chimeras can be used to uptake soluble and/or cell membrane-bound cargo selectively into CD206^+^ cells (**Fig. 1b, c**). This antibody-based ligand screen led to the identification of a new class of small molecule ligands for CD206. Moreover, this approach is modular and agnostic to the choice of the antibody employed, enabling virtually any antibody to be rapidly transformed into a TICTAC, including those targeting immune-checkpoint proteins. We demonstrate that TICTACs can facilitate robust depletion of immune checkpoint receptors from the surface of CD206^hi^, but not CD206^lo^ macrophages. This approach may lead to a new class of immune therapies with reduced risk of systemic immune activation and associated toxicities.

## Results

### Design and synthesis of multivalent CD206-targeting chimeras

Prior works targeting CD206 largely employs mannose-functionalized glycopolymers that use avidity to compensate for the low affinity of monomeric mannose^16–22^. However, the heterogeneous nature of these glycopolymers render reproducible synthesis and characterization of the resulting antibody-conjugates challenging. Therefore, we sought to develop alternative low molecular weight homogeneous ligands that bind CD206. A recent glycan array study identified L-fucose as a promising candidate: >60% of the top 20 binders were fucosylated, and fucose-containing glycans were structurally simpler than those containing mannose^23^. We also explored sulfated glycans such as 3-SO_4_ galactose, which bind the N-terminal R-type carbohydrate recognition domain (R-type CRD) instead of the C-type lectin-like domains (CTLDs) commonly targeted by mannose (**Fig. 1a**). The R-type CRD was first shown to bind 4-SO_4_-GalNAc^24^, and subsequent studies demonstrated binding to 3-SO_4_-Gal/GalNAc^25^. Therefore, based on the known carbohydrate specificity of CD206, we rationally designed and synthesized a focused set of propargyl-, azide-, or aminooxy-functionalized mannose, fucose, and 3-SO4 galactose mono-, di-, and trisaccharides (**Extended Data Fig. 1**). Antibody-glycan conjugates were generated in a two-step sequence, where goat-anti-rabbit secondary antibodies were first modified on lysine residues with linkers displaying alkynes, azides, or aldehydes in various valencies and orientations, then conjugated to the carbohydrate ligands via Cu-catalyzed or strain-promoted click reactions, or oxime ligations (**Fig. 1b**). Characterization of the antibody-glycan conjugates by MALDI-MS analysis revealed average ligand-to-antibody ratios between 10 and 16 (**Supplementary Fig. 1**).

We next sought to identify a suitable *in vitro* macrophage model system. Phorbol 12-myristate 13-acetate (PMA)-differentiated THP-1 or U-937 cells are frequently utilized as human macrophage surrogates, however, polarization with IL-4, IL-10, or IL-13 all failed to induce expression of CD206^26^. We thus turned to the murine macrophage cell line RAW264.7, another extensively used model for studying macrophage biology^27^. Gratifyingly, addition of 25 ng/mL IL-4 over 24 hours resulted in robust upregulation of CD206 (M2-polarized) in comparison to unpolarized cells (**Fig. 1c**).

### Trivalent TICTACs drive CD206-dependent internalization in macrophages

We examined the ability of the antibody-glycan conjugates to internalize an extracellular target, rabbit IgG-488, via CD206, leading to the identification of trivalent TICTACs bearing dendritic fucose, 3-SO_4_ galactose, or mannose ligands (**1a-c**, respectively) (**Fig. 1d, Extended Data Fig. 2**). Degradation studies comparing monovalent or multivalent ligand antibody conjugates revealed that tris-ligand presentation yielded more efficient CD206-mediated uptake (**Supplementary Fig. S2**). Specifically, trivalent dendron constructs (t1a, t1b, and t2b) consistently outperformed mono-, bi-, and pentavalent scaffolds in direct comparisons (**Supplementary Fig. S2b**), and the PEG5-based linker length was identified as optimal for CD206-mediated uptake through empirical screening of the panel. M2-polarized RAW264.7 cells were incubated with rabbit IgG-488 and goat-anti-rabbit (control) or goat-anti-rabbit TICTACs **1a-c** for 3 hours, then analyzed by flow cytometry for 488 fluorescence (**Fig. 1e**). Because of the broad expression profile of CD206, cells were co-stained with an anti-CD206 antibody and gated for CD206^hi^ and CD206^lo^ populations. In the CD206^hi^ population, treatment with tris-3-SO_4_ Gal TICTAC **1b** resulted in a 6-fold increase in intracellular fluorescence relative to the unconjugated parent antibody, while tris-Fuc TICTAC **1a** and tris-Man TICTAC **1c** gave 2.6-fold and 2-fold increases, respectively (**Fig. 1f**). Direct triazole linkage at the anomeric position of mannose (**1c\***) completely ablated internalizing activity. In the CD206^lo^ population, **1a**-**d** did not result in significant increases in intracellular fluorescence (**Fig. 1f**). Additionally, examination of Fuc and Man ligand conjugates revealed that the tris-Fuc ligand yielded more efficient CD206-mediated uptake compared to the analogous tris-Man (**Supplementary Fig. S2**).

We then evaluated whether TICTAC-mediated uptake was generalizable across other macrophage cell lines. Internalization of fluorophore-conjugated rabbit-IgG was observed for M2-polarized BMA3.1A7^28^ and J774A.1 cell lines using TICTACs **1a**-**c** (**Fig. 1g** *left*, **Supplementary Fig. S3**), with **1a** and **1b** showing superior uptake. The M2-polarized microglial cell line BV-2 was also treated with the unconjugated parent antibody, TICTACs **1a**, **1b**, or a tris-Gal analogue (negative control), resulting in internalization of **1a** and **1b**, but not tris-Gal (**Fig. 1g** *right*). Finally, to validate the mechanism of uptake, we generated a CD206-knockout RAW264.7 cell line and subjected it to M2-polarization conditions before treatment with fluorophore-conjugated rabbit IgG and unconjugated goat-anti-rabbit control, **1a**, or **1b** (**Fig. 1h**). Uptake was abolished in the knockout cells, while preserved in cells that were treated with Cas9 without sgRNA (mock control).

### Small molecule TICTAC ligands are internalized by M2-polarized human macrophages

We next asked whether the conjugated tris-fucose ligand in **1a** could function independently as a small molecule CD206 ligand. We synthesized tris-Fuc 647 (**2**), wherein the tris-fucose ligand was conjugated to a 647-dye in a 1:1 stoichiometry (**Fig. 2a, Supplementary Note 2**). Polarized RAW264.7 cells were treated with various concentrations of the unconjugated dye or compound **2** for 2 h, and fluorescence was analyzed by flow cytometry (**Fig. 2b**). Uptake was dependent on the concentration of **2**, and significant differences were observed between **2** and the unconjugated 647-dye control at concentrations as low as 0.1 nM. This result was confirmed by confocal microscopy, where treatment with **2** resulted in a high 647 signal that concentrated in certain cells over others, mirroring the broad expression profile of CD206 (**Fig. 2c**). Next, we tested whether **2** is internalized by primary human macrophages. Upregulation of CD206 was first confirmed upon polarizing the macrophages with IL-4/IL-10 and TGF-β over several days^29^ (**Fig. 2d**). M2-polarized and non-polarized macrophages were subjected to the unconjugated dye or **2** at various concentrations, and intracellular fluorescence was measured by flow cytometry (**Fig. 2e**). In addition to a significant increase in fluorescence observed by treatment with **2** compared to the unconjugated control, we observed a notable preference for M2-polarized over nonpolarized cells. CD206-knockout RAW264.7 cells were unable to internalize compound **2**, confirming the selectivity of the ligand for CD206 (**Fig. 2f**). Binding of **2** to CD206 was also analyzed by surface plasmon resonance (SPR), which revealed an extremely slow off rate (<2.4 x 10^−4^ min^−1^) that likely drives internalization even at sub-nanomolar concentrations (**Extended Data Fig. 3**).

**Figure 2.**
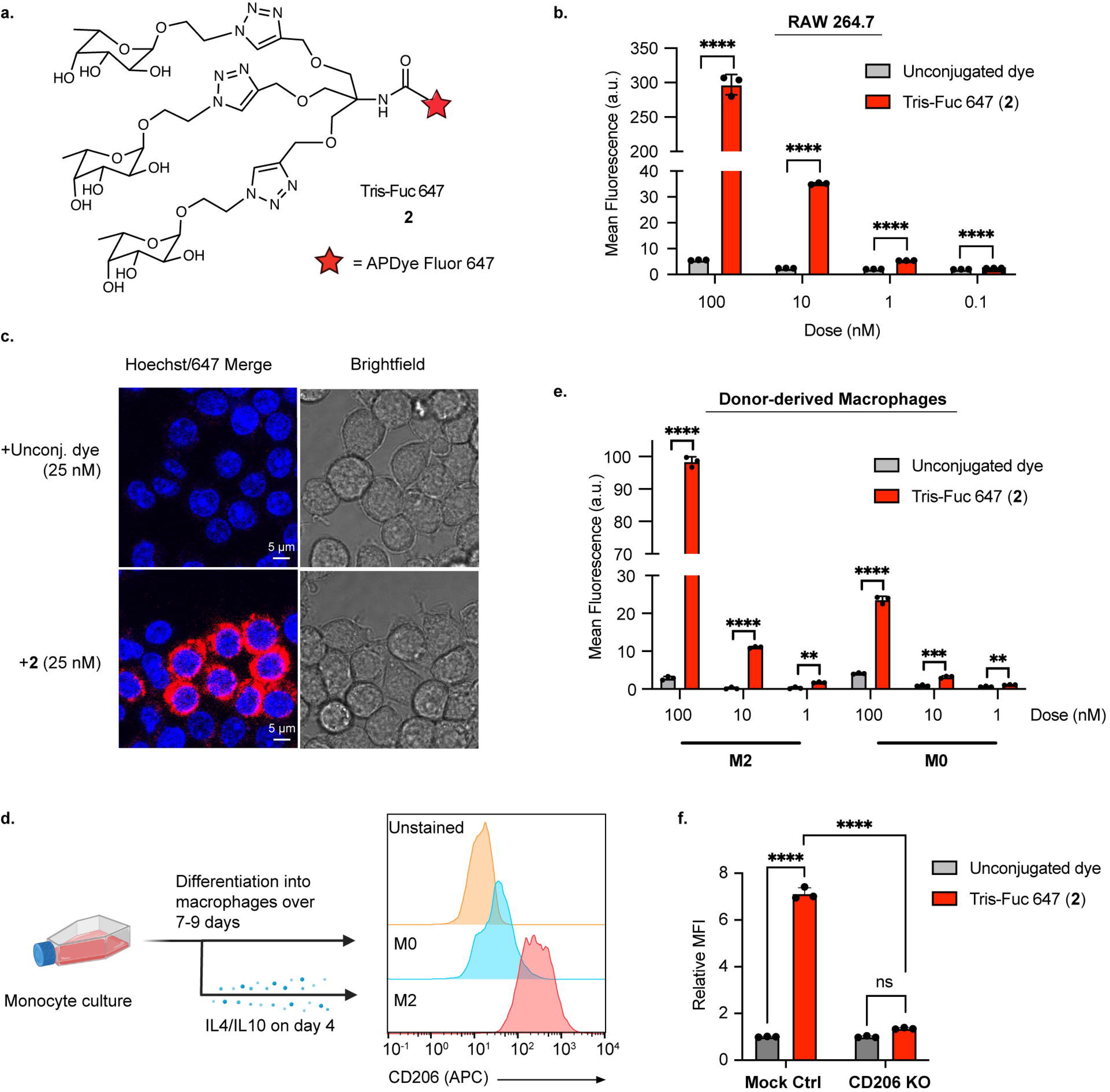
Tris-fucose ligand functions independently as a small molecule CD206 binder. **a,** Tris-fucose ligand was independently synthesized and conjugated to a 647 dye to generate tris-Fuc-647 **2**. **b**, Changes in 647 mean fluorescence intensity (MFI) in M2-polarized RAW264.7 cells (gated for CD206^hi^ population) upon treatment with unconjugated 647 dye or **2** at varying concentrations for 2 h at 37 °C. Data are meanL±Ls.d. of *n*L=L3 biological replicates for each condition/dose. **c,** Confocal microscopy imaging of M2-polarized RAW264.7 cells treated for 2 h with 25 nM **2** or unconjugated 647 dye. Data are representative of 2 independent experiments with similar results. **d,** Monocytes were isolated from human PBMCs and differentiated into macrophages either in the absence or presence of IL-4 (20 ng/mL) and IL-10 (50 ng/mL). CD206 is upregulated in M2-polarized primary human macrophages. **e,** Changes in 647 MFI in M2-polarized vs. non-polarized human macrophages upon treatment with unconjugated 647 dye or **2** at varying concentrations for 2 h at 37 °C. Data are meanL±Ls.d. of *n*L=L3 independent experiments using macrophages derived from a single human donor. **f,** Knocking out CD206 ablates uptake of **2** in RAW264.7 cells. Data are meanL±Ls.d. of *n*L=L3 biological replicates. For **b** and **e**, *P* values were determined by parametric two-tailed *t*-tests and by two-tailed unpaired t-test for **f**. Statistical significance was defined as *P*<0.05, and the asterisks ** indicates *P*<0.01, *** indicates *P*<0.001, and **** indicates *P*<0.0001.

### Non-blocking TICTACs mediate CD206-dependent removal of soluble cytokines

In addition to TAM-specific delivery of exogenous small molecule or protein-based cargo, we investigated whether TICTACs could clear endogenous soluble targets such as cytokines selectively. To test this strategy, we chose to target IL-4, a driver for M2 polarization in TAMs and immune suppression within the tumor microenvironment^30^. We generated TICTACs **3a** and **3b** through conjugation of tris-Fuc and tris-3-SO_4_ Gal ligands respectively with a previously characterized non-neutralizing anti-IL4 antibody (clone BVD6-24G2)^31^ and subjected polarized RAW264.7 cells to 488-labeled murine IL4 and unconjugated anti-IL4 antibody, **3a**, or **3b** (**Extended Data Fig. 4a**). In CD206^hi^ populations, **3a** and **3b** induced >10-fold increase in intracellular 488 fluorescence relative to the unconjugated antibody, whereas the change in CD206^lo^ populations was less than 2-fold (**Extended Data Fig. 4b**). These results suggest the potential for non-blocking TICTACs to be employed for tumor-specific removal of soluble targets.

### TICTACs mediate robust, IL-4 dependent depletion of membrane protein CD54

Having demonstrated that TICTACs can efficiently internalize soluble extracellular cargo, we next investigated whether they could mediate depletion of membrane proteins in M2-polarized macrophages. For proof-of-concept, we chose CD54 (ICAM-1) as a target due to its robust expression in several macrophage cell lines. TICTACs were constructed using a commercially available primary antibody against CD54 (clone 3E2) following a similar scheme to **1a** and **1b** to generate tris-Fuc conjugated **4a** and tris-3-SO_4_ Gal conjugated **4b** (**Fig. 3a, Supplementary Fig. S4**). No significant change in chemical stability was observed between the unconjugated antibody and the Tris-conjugated TICTAC (**Supplementary Fig. S5**). We then treated RAW264.7 cells with **4a** and **4b**, and measured surface levels of CD54 by flow cytometry using an orthogonal detection antibody. TICTAC **4a** resulted in >80% depletion of surface CD54 in CD206^hi^ RAW264.7 cells, whereas TICTAC **4b** resulted in >60% depletion. In CD206^lo^ cells, this effect is greatly attenuated, with **4a** and **4b** treatment resulting in 30% and 15% depletion of surface CD54, respectively (**Fig. 3b**, **Supplementary Fig. S6**). We note that relative surface CD54 was calculated by normalizing the MFI of the detection antibody to the unconjugated antibody control within each gated population (CD206^hi^ or CD206^lo^) independently. CD54 is robustly expressed in both M0 and M2 macrophages, so differences in basal expression do not confound the analysis. TICTAC activity was also inhibited in CD206-knockout RAW264.7 cells, further providing support for a CD206-mediated mechanism of action (**Supplementary Fig. S7**). We also evaluated the activity of **4a** in BMA3.1A7 cells and J774A.1 cell lines. We observed >60% depletion of surface CD54 in CD206^hi^ BMA3.1A7 cells, while <30% depletion was observed in CD206^lo^ cells (**Fig. 3c**). J774A.1 cells also exhibited comparable reduction in cell surface CD54, which was dependent on expression levels of CD206 (**Supplementary Fig. S8a**). Moreover, we found that cell surface CD206 levels remain unaffected by presence of unconjugated anti-CD54 antibody or tris-Fuc conjugated **4a** TICTAC (**Extended Data Fig. 5a, b**), and that TICTAC treatment did not affect the cell surface levels of other surface markers (**Extended Data Fig. 5c**). Visualization of surface CD54 by confocal microscopy following TICTAC treatment showed significantly reduced membrane CD54, consistent with our flow cytometry observations (**Fig. 3d**). Reduction of cell surface CD54 on CD206^hi^ RAW264.7 cells was concentration dependent, where **4a** demonstrated a detectable decrease in surface CD54 at concentrations as low as 0.1 nM. Maximum activity of **4a** was reached at 5 nM, where the same degree of depletion was maintained at higher concentrations up to 100 nM with no observable “hook effect”^32^ (**Fig. 3e**). Conjugates **4a** and **4b** depleted cell-surface CD54 over 48 h, where >50% reduction was observed for **4a** after 3 h, which further increased to >80% removal after 24 h (**Fig. 3f, Supplementary Fig. S9**). Depletion of cell-surface CD54 mediated by **4b** was markedly slower than by **4a**. Finally, because CD206 expression is dependent on the cytokine IL-4, we reasoned that TICTAC activity must also be dependent on IL-4. We set up two parallel experiments where RAW264.7 or J774A.1 cells were either polarized with IL-4 or not prior to treatment with **4a**. We observed **4a**-mediated depletion of cell-surface CD54 that was dependent on the presence of IL-4, demonstrating that TICTACs can be “turned on” with IL-4 (**Fig. 3g**, **Supplementary Fig. S8B**).

**Figure 3.**
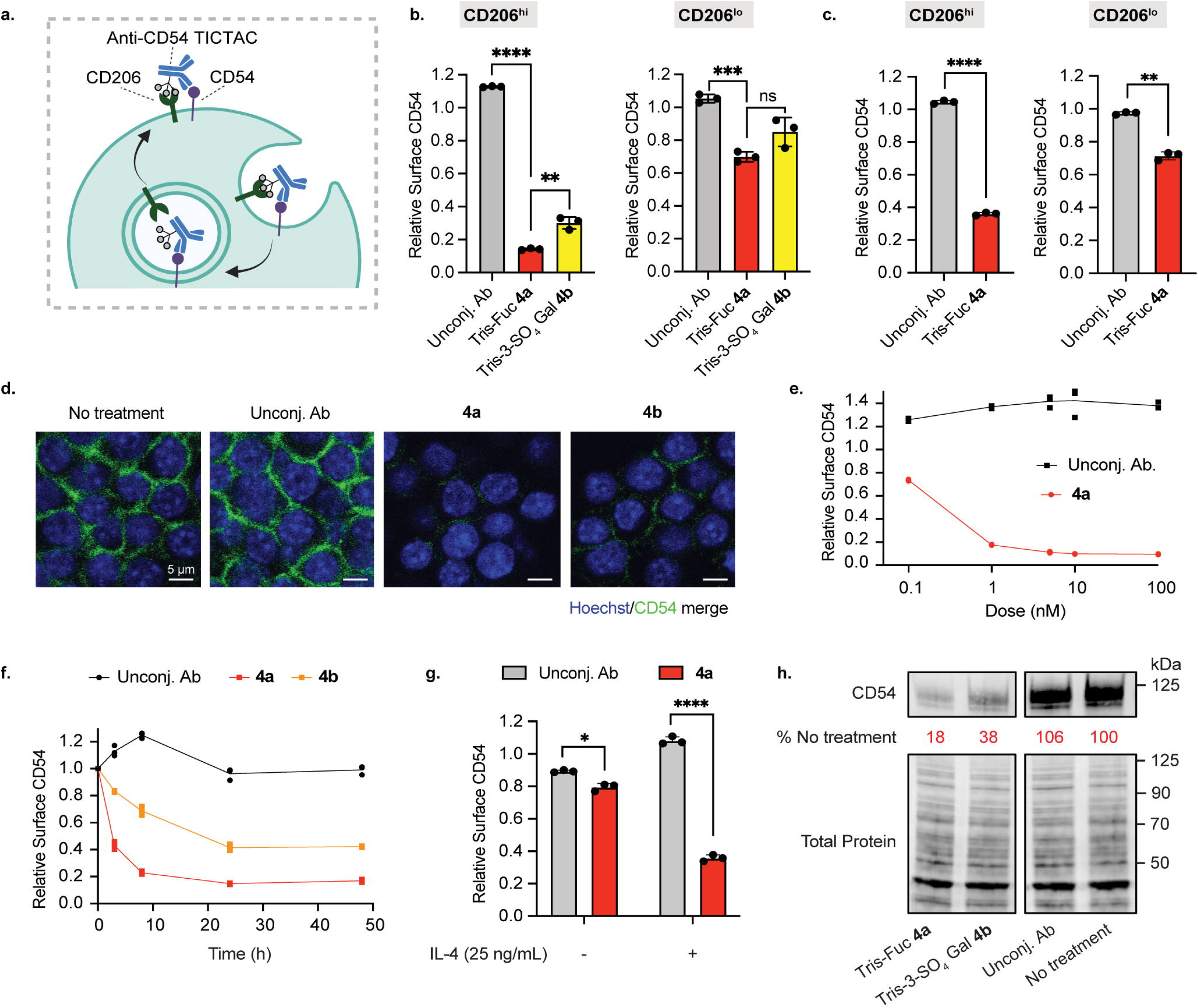
Anti-CD54 TICTAC depletes cell-surface CD54 in a CD206-dependent manner. **a**, Proposed mechanism for the removal of cell-surface CD54 mediated by TICTACs. **b,** Depletion of cell-surface CD54 in M2-polarized RAW264.7 cells determined by live cell flow cytometry following 24 h of treatment with 25 nM unconjugated anti-CD54 antibody or conjugates. Data are meanL±Ls.d. of *n*L=L3 biological replicates. **c,** Depletion of cell-surface CD54 in M2-polarized BMA3.1A7 cells. Data are meanL±Ls.d. of *n*L=L3 biological replicates. **d,** Visualization of cell-surface CD54 by confocal microscopy after 25 nM TICTAC treatments for 24 h. Data are representative of 2 independent experiments with similar results. **e,** Dose-response curve for cell-surface CD54 removal in M2-polarized RAW264.7 cells following treatment with unconjugated antibody or **4a** at 0.1 nM, 1 nM, 5 nM, 10 nM, and 100 nM for 24 h. **f,** Time-course of CD54 downmodulation in M2-polarized RAW264.7 cells incubated with 25 nM unconjugated antibody or conjugates at 3 h, 8 h, 24 h, and 48 h. **g,** Depletion of cell-surface CD54 in RAW264.7 cells that have been polarized with IL-4 (25 ng/mL) or not following treatment with 25 nM unconjugated antibody or **4a**. Data are meanL±Ls.d. of *n*L=L3 biological replicates. **h,** Degradation of CD54 in M2-polarized bone marrow-derived macrophages harvested from C57BL/6 mice after treatment with **4a** and **4b** for 24 h. Data are representative of three independent experiments. For **b** and **c**, *P* values were determined by one-way ANOVA followed by Tukey’s post-hoc test for multiple comparisons. For **g**, *P* values were determined by Welch’s two-tailed *t*-tests. Statistical significance was defined as *P*<0.05, and the asterisks * indicates *P*<0.05, ** indicates *P*<0.01, *** indicates *P*<0.001, and **** indicates *P*<0.0001.

To determine whether internalized CD54 was subsequently degraded, we measured total CD54 levels in RAW264.7 cells following treatment with **4a** or **4b** by Western blot (**Extended Data Fig. 6a**). Surprisingly, TICTAC treatment induced only small changes in total protein levels (20% lower than no treatment) despite high levels of cell-surface CD54 depletion as observed by flow cytometry. We next evaluated TICTAC activity in primary murine bone marrow-derived macrophages (BMDMs), which were isolated from C57BL/6 or BALB/c WT mice. BMDMs were polarized with 25 ng/mL murine IL-4 prior to treatment with TICTACs or unconjugated antibody control for 24 h. TICTAC treatment resulted in >80% degradation of total CD54 for tris-Fuc conjugate **4a** and >60% degradation for tris-3-SO_4_ Gal conjugate **4b** relative to no treatment or the unconjugated antibody control (**Fig. 3h, Extended Data Fig. 6b**). Additionally, we measured cell surface CD54 and CD206 levels by flow cytometry (**Supplementary Fig. 10**). We found that while CD54 levels were reduced for TICTAC-treated BMDMs, compared to the vehicle or unconjugated anti-CD54 antibody, cell-surface CD206 levels remained unchanged. It is critical to note this stark contrast in the degradative ability of the TICTACs between immortalized cell lines (such as RAW264.7) and primary macrophages. Previous mass spectrometry based proteomics analyses of both cell types revealed prominent differences in protein profiles associated with polarization, phagocytosis, and innate immune responses^33,34^. CD206, along with proteins regulating early to late endosome transport and phagosome maturation, were markedly attenuated in macrophage cell lines, which may explain why TICTACs induce more potent degradation in primary BMDMs than RAW264.7 cells.

### TICTACS selectively degrade the SIRP***α*** checkpoint in M2 macrophages

We next sought to determine whether TICTACs could provide an alternative form of immune checkpoint blockade that was specific to M2-polarized macrophages, which are known to drive immune suppression in the tumor microenvironment. In a canonical checkpoint blockade, the antibody drug systemically blocks the immune checkpoint protein (ICP) and activates immune cells in healthy and cancerous tissue alike, leading to irAEs. We hypothesized that by generating TICTACs from *non-blocking* antibodies that bind but do not inhibit the ICP, we could selectively remove the ICP from TAMs while mitigating off-target effects (**Fig. 4a**). We targeted signal regulatory protein alpha (SIRPα), which is highly expressed on myeloid-derived immune cells and potently engages the “don’t eat me” ligand CD47^35–39^. Monoclonal antibodies against both SIRPα and CD47 are currently being evaluated as cancer immune therapies^37,40,41^. Blocking antibody clone 119 and non-blocking antibody clones 136 and 3 were recently developed as pan-allelic, high affinity SIRPα binders^42^ (**Fig. 4b**).

**Figure 4.**
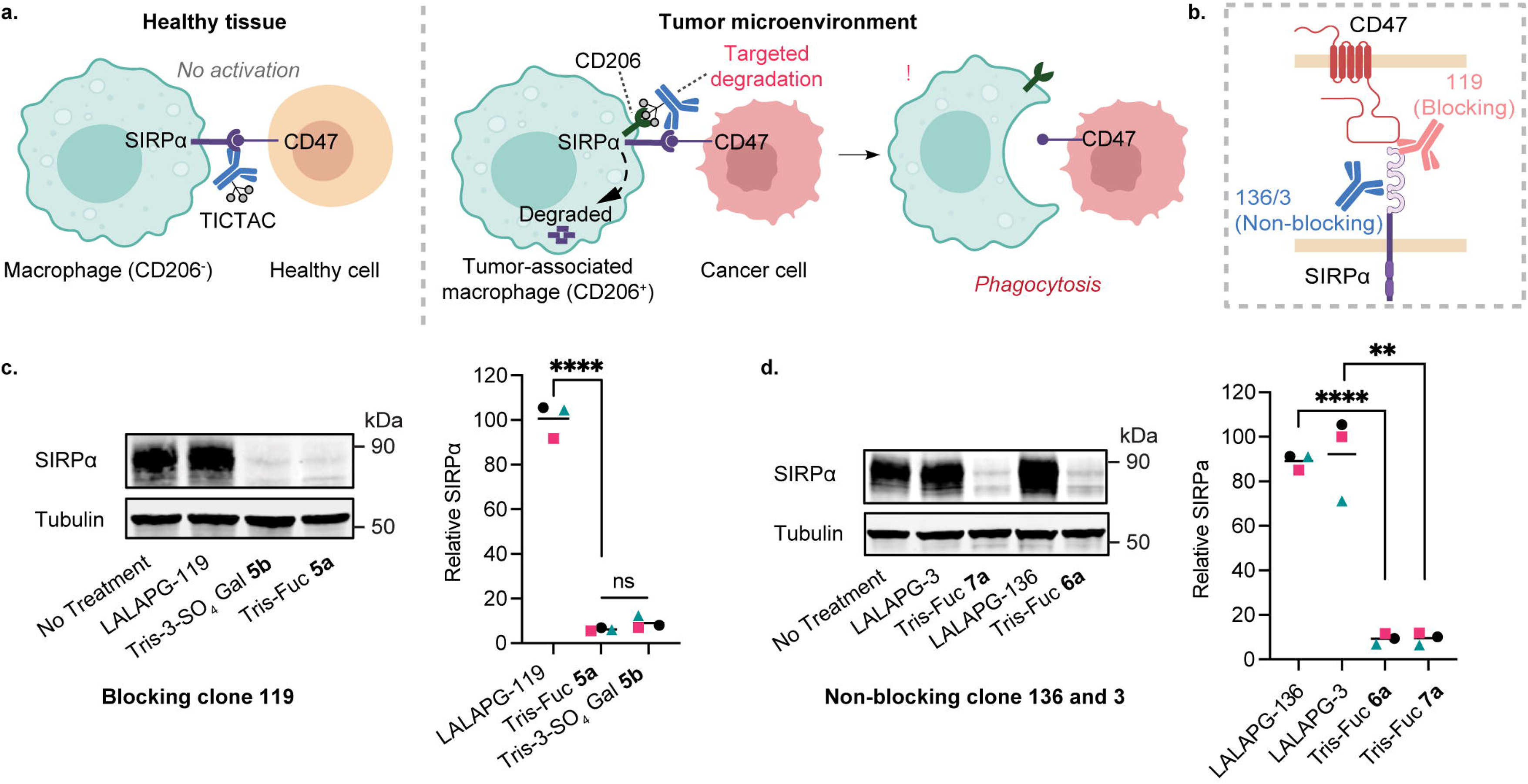
Anti-SIRPα TICTACs degrade immune checkpoint protein SIPRα regardless of their ability to block CD47. **a,** Non-blocking TICTACs can degrade ICPs in CD206^hi^ TAMs (*left*) but have no function-ablating activity in healthy tissue (*right*). **b,** While antibody clone 119 blocks the CD47-binding site, antibodies 136 and 3 bind a distal, non-blocking site. **c,** Tris-Fuc and 3-SO_4_-Gal TICTACs generated from blocking clone 119 (**5a**, **5b**) induce degradation of SIRPα in human M2-polarized macrophages following treatment at 25 nM for 24 h. Data are mean of *n*L=L3 human donors, with each point representing mean of 3 independent experiments per donor. **d,** Tris-Fuc TICTACs **6a** and **7a** generated from non-blocking clones 136 and 3 respectively induce degradation of SIRPα in human M2-polarized macrophages following treatment at 25 nM for 24 h. Data are mean of *n*L=L3 human donors, with each point representing mean of 3 independent experiments per donor. For **c** and **d**, *P* values were determined by one-way ANOVA followed by Tukey’s post-hoc test for multiple comparisons. Statistical significance was defined as *P*<0.05, and the asterisks * indicates *P*<0.05, ** indicates *P*<0.01, *** indicates *P*<0.001, and **** indicates *P*<0.0001.

We first generated LALAPG-119, LALAPG-136, and LALAPG-3 using the corresponding Fab fragments and human IgG1 containing Fc-silencing mutations L234A/L235A/P329G (‘LALAPG’) to mitigate competing Fc receptor (FcR) binding^43^. TICTACs were assembled following a similar scheme to **1a** and **1b**, generating tris-Fuc and tris-3-SO_4_-Gal functionalized LALAPG-119 (**5a** and **5b** respectively), tris-Fuc functionalized LALAPG-136 (**6a**), and tris-Fuc functionalized LALAPG-3 (**7a**) (**Supplementary Fig. S11–13**). M2-polarized primary human macrophages were subjected to 25 nM unconjugated LALAPG-119, **5a**, or **5b** for 24 h, and total SIRPα levels were quantified by Western blot analysis (**Fig. 4c**). While the unconjugated antibody had no effect on SIRPα levels, TICTAC treatment resulted in almost complete (>90%) degradation of SIRPα across three human donors. We next asked whether TICTACs that bind different epitopes of the target protein could also induce selective degradation. M2-polarized macrophages were treated with non-blocking LALAPG-136, 3, or the corresponding tris-Fuc TICTACs **6a** and **7a**. Upon assessing SIRPα levels by Western blot, non-blocking TICTACs were equally as effective as their blocking counterpart, again resulting in >90% degradation across three independent donors (**Fig. 4d**). These results demonstrate that TICTACs enable decoupling of receptor binding from the inhibitory effect (*via* degradation), giving rise to highly TAM-specific activity. Further, we observed high levels of SIRPα degradation (>70% relative to no treatment control) when we treated M2-polarized primary murine BMDMs with **6a** (25 nM for 24 h), indicating excellent cross-reactivity in mice (**Extended Data Fig. 7**).

We also used quantitative mass spectrometry-based proteomics to measure proteome-wide changes in human macrophages (across 3 independent donors) treated with LALAPG-119 or **5a** (**Fig. 5a**). While no significant changes were observed upon treatment with LALAPG-119, **5a** promoted a marked reduction in SIRPα levels. Moreover, TICTACs exert high target precision: downmodulation was highly localized to SIRPα across the entire proteome. We next wanted to determine the selectivity of anti-SIRPα TICTAC activity for M2-polarized macrophages over non-polarized macrophages and other immune cells. M2- (CD206^hi^) and M0-polarized (CD206^lo^) macrophages were generated from the same donor and treated with LALAPG-119 or **5a**. While M2-macrophages again exhibited substantial TICTAC-induced degradation (90% degradation relative to no treatment), this effect was greatly attenuated in M0-macrophages (30% degradation relative to no treatment) which expressed basal levels of CD206 (**Extended Data Fig. 8**). Signal regulatory protein gamma (SIRPγ) is another member of the SIRP family that is uniquely expressed by human T-cells. While its function is less well-characterized, its engagement with CD47 on antigen presenting cells (APCs) enhances antigen-specific T-cell proliferation and facilitates endothelial migration of T-cells *in vitro*^44,45^. Because many antibodies targeting SIRPα are cross-reactive with SIRPγ, we wanted to interrogate whether TICTACs exhibited any off-target degradation of SIRPγ expressed in human T-cells. PBMCs isolated from three independent donors were subjected to LALAPG-119, LALAPG-136, or the corresponding TICTACs **5a** and **6a**. After 24 h, the cells were analyzed by flow cytometry, where T-cells were identified as the CD3^+^ population (**Supplementary Fig. S14**). Upon quantifying cell-surface SIRPγ levels, we found that TICTACs induced no appreciable downmodulation of SIRPγ relative to the unconjugated antibody. These data indicate TICTAC-mediated degradation is highly specific to M2-polarized macrophages, preserving target protein expression on non-polarized macrophages and other immune cells.

**Figure 5.**
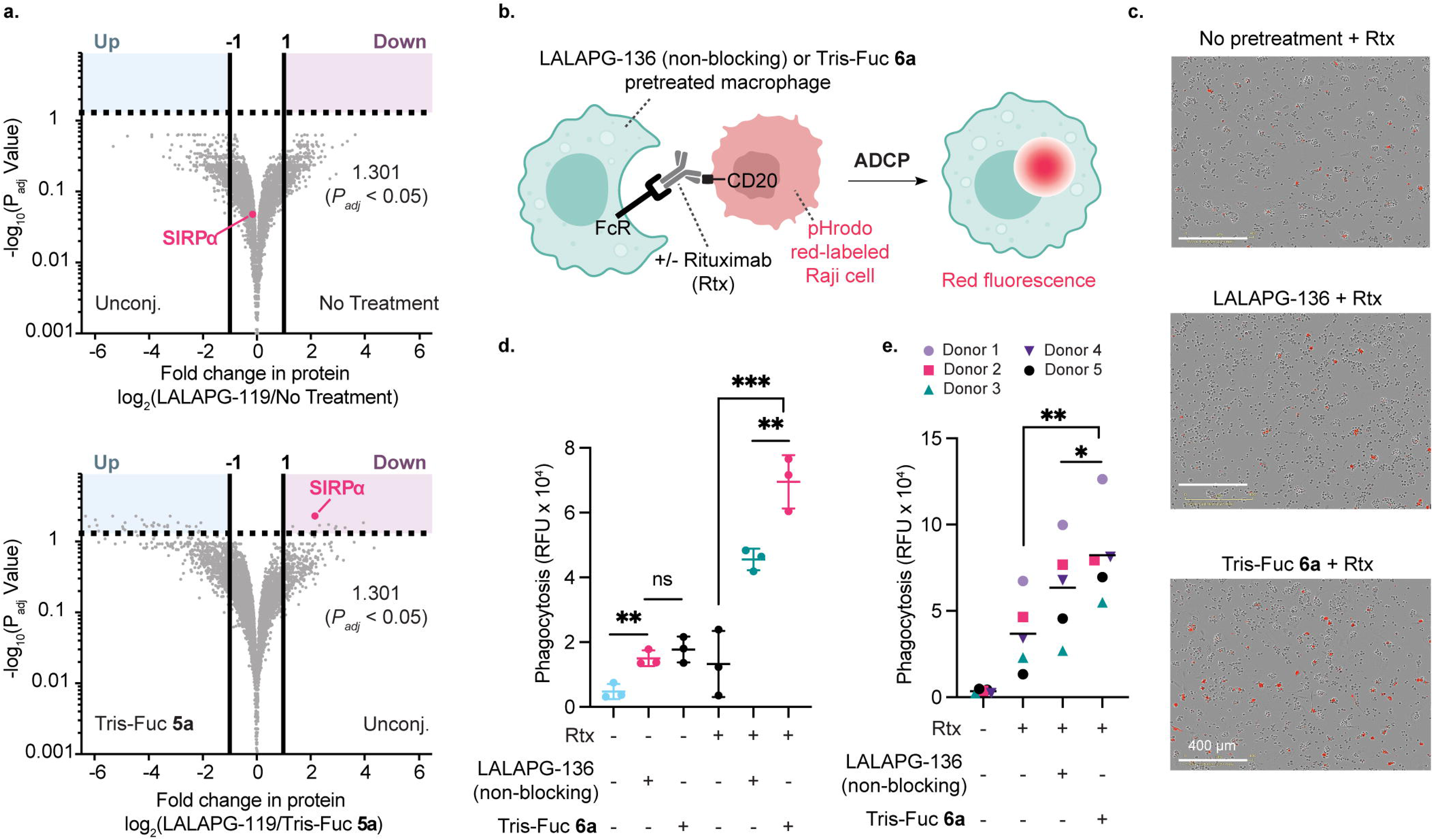
Anti-SIRPα TICTACs are specific for SIRPα and significantly enhance phagocytosis in TAMs. **a,** Fold change in the abundance of 5,091 proteins detected by quantitative proteomics analysis after 24 h treatment with either unconjugated LALAPG-119 or **5a**. The no treatment sample represents a PBS control. SIRPα (highlighted in fuchsia) is downregulated in **5a**-treated macrophages relative to unconjugated 119. An adjusted *P*-value (*P*_adj_) threshold of 0.05 and a log_2_ fold change threshold of 1.0 were set to identify significant changes to protein abundance. Data are the mean of three independent donors. **b,** Donor-derived macrophages pretreated with either no antibody (PBS control), LALAPG-136, or Tris-Fuc **6a** are co-cultured with pHrodo red-labeled Raji cells in the presence or absence of Rituximab (Rtx). The resulting ADCP was quantified by total red fluorescence signal. **c,** Phase contrast and fluorescence microscopy images of human macrophages/Raji co-cultures treated with rituximab at 5 µg/mL. Macrophages were pre-treated with 25 nM unconjugated LALAPG-136 or **6a** for 24 h prior to co-culture. Images are representative of 3 independent experiments from a single donor. Scale bars represent 400 µm. **d,** Phagocytosis of Raji cells by M2-polarized human macrophages treated with LALAPG-136 or **6a** in the presence or absence of rituximab. Data are mean of *n*L=L3 independent experiments using macrophages derived from a single human donor. *P* values were determined by unpaired two-tailed *t*-tests. **e,** Phagocytosis of Raji cells by M2-polarized human macrophages treated with LALAPG-136 or **6a** in the presence or absence of rituximab. Data are mean of *n*L=L5 human donors, with each point representing mean of 3 independent experiments per donor. *P* values were determined by paired two-tailed *t*-tests. For **d**, and **e**, statistical significance was defined as *P*<0.05, and the asterisks * indicates *P*<0.05, ** indicates *P*<0.01, *** indicates *P*<0.001, and **** indicates *P*<0.0001.

### SIRP***α*** degradation enhances macrophage-mediated tumor phagocytosis

We further asked whether TICTACs could have a functional anti-tumor effect on M2-polarized macrophage activity. Antibody-dependent cellular phagocytosis (ADCP) and antibody-dependent cellular cytotoxicity (ADCC) are tumoricidal processes resulting from co-engagement of FcRs on immune cells and antigens on the cancer cell surface. Because inhibitory checkpoint receptors such as SIRPα transmit “don’t eat me” signals that potently block ADCP, we hypothesized that TICTAC-mediated degradation of SIRPα would enhance phagocytosis in the presence of therapeutic tumor-targeting antibodies. To test this, we first treated M2-polarized human macrophages with LALAPG-136 or **6a** for 48 h, then co-cultured them with Raji cells that were labeled with pHrodo red, a pH-sensitive dye that fluoresces red in acidic phagosomes (**Fig. 5b**). Fluorescence microscopy images were acquired at 1 h intervals, and phagocytosis was quantified as total integrated red fluorescence normalized by cellular confluence in each well (**Fig. 5c, d**). Maximum phagocytosis was observed after 4 h of co-culture in the presence of rituximab, a CD20-targeting therapeutic antibody. A somewhat unexpected result was that pre-treatment with the unconjugated LALAPG-136 control induced an increase in ADCP relative to the vehicle control, despite its lack of blocking activity (**Fig. 5d**). This background effect aligns with previous reports showing that clone 136 can modestly enhance ADCP in donor-derived macrophages, albeit much less potently than blocking clone 119^42^. Consistent with those studies, we do not attribute this background activity to the downregulation of SIRPα, which was confirmed by Western blot (**Fig. 4d**), but rather to subtle alterations in tonic SIRPα signaling that occur independently of CD47 engagement. Importantly, TICTAC treatment further enhanced phagocytosis beyond both the vehicle and unconjugated LALAPG-136 controls, and this effect was consistent across macrophages from 5 donors (**Fig. 5e**). Moreover, we conducted additional experiments to address the possibility that while cell-surface and total levels of CD206 are unchanged, there may still be discrepancies in CD206 trafficking function. It is known that one of the key biological functions of CD206 is collagen internalization, which represents a major mechanism of collagen turnover in mice^46,47^. Given that CD206 binds to collagen via its fibronectin type II (FNII) domain, we adapted a published protocol for a quantitative *in vitro* collagen uptake assay^48^ (see Methods) to examine whether TICTAC treatment alters the ability of M2 macrophages to endocytose fluorescently labeled collagen (**Extended Data Fig. 9 and Supplementary Fig. S15**). We found that neither tris-Fuc nor tris-SO_4_-Gal antibody conjugates (nor the tris-Fuc small molecule) significantly affects collagen uptake. Taken together, these results show that TICTACs can have a functional effect on antibody effector functions such as ADCP in M2-polarized macrophages while preserving the native biological function of CD206.

### Anti-SIRP***α*** TICTACs reduce tumor burden and metastasis in vivo

Finally, to address the *in vivo* efficacy of TICTACs, we developed a humanized NSG mouse model, which recapitulates human donor-specific immune phenotypes and enables a xenograft of human cancer cells (**Fig. 6**). Briefly, Raji cells (B cell lymphoma) were implanted subcutaneously into immunodeficient NSG mice and the tumor was allowed to develop for nine days. On day nine, each mouse received an intraperitoneal dose of human donor-derived PBMCs and TAMs at a 10:1 PBMC:TAM ratio. TAMs were generated by isolating monocytes from PBMCs and polarizing them *ex vivo* into M2 macrophages using IL-4, IL-10, and TGF-β prior to adoptive transfer (see Methods). Beginning on day fourteen, mice were injected with either rituximab (anti-CD20 antibody) or a co-treatment of rituximab and anti-SIRPα Tris-Fuc **6a** TICTAC (**Fig. 6a**). While treatment with rituximab failed to slow tumor progression, co-treatment with anti-SIRPα TICTAC resulted in a significant reduction in tumor burden (**Fig. 6b-e**). Concordantly, immunohistochemistry (IHC) staining of excised xenografts showed enhanced apoptosis (cleaved caspase-3), lower proliferation (Ki-67), reduced microvascular density (CD31), reduced SIRPα expression, and reduced lung metastasis in TICTAC-treated mice (**Extended Data Fig. 10a and Fig. 6g-k**). Notably, the SIRPα signal in these tumor sections is attributed to the adoptively transferred human macrophages. Raji cells do not express SIRPα, which is a myeloid lineage-restricted receptor^49–51^, and the anti-SIRPα antibody used for the IHC is specific for human SIRPα and does not cross-react with the murine ortholog. Lastly, we found that TICTAC treatment results in reduced metastasis in the blood vessels and alveoli of mice lung compared to the rituximab-only group (**Extended Data Fig. 10b**). Together, these data demonstrate the therapeutic efficacy of TICTACs *in vivo*.

**Figure 6.**
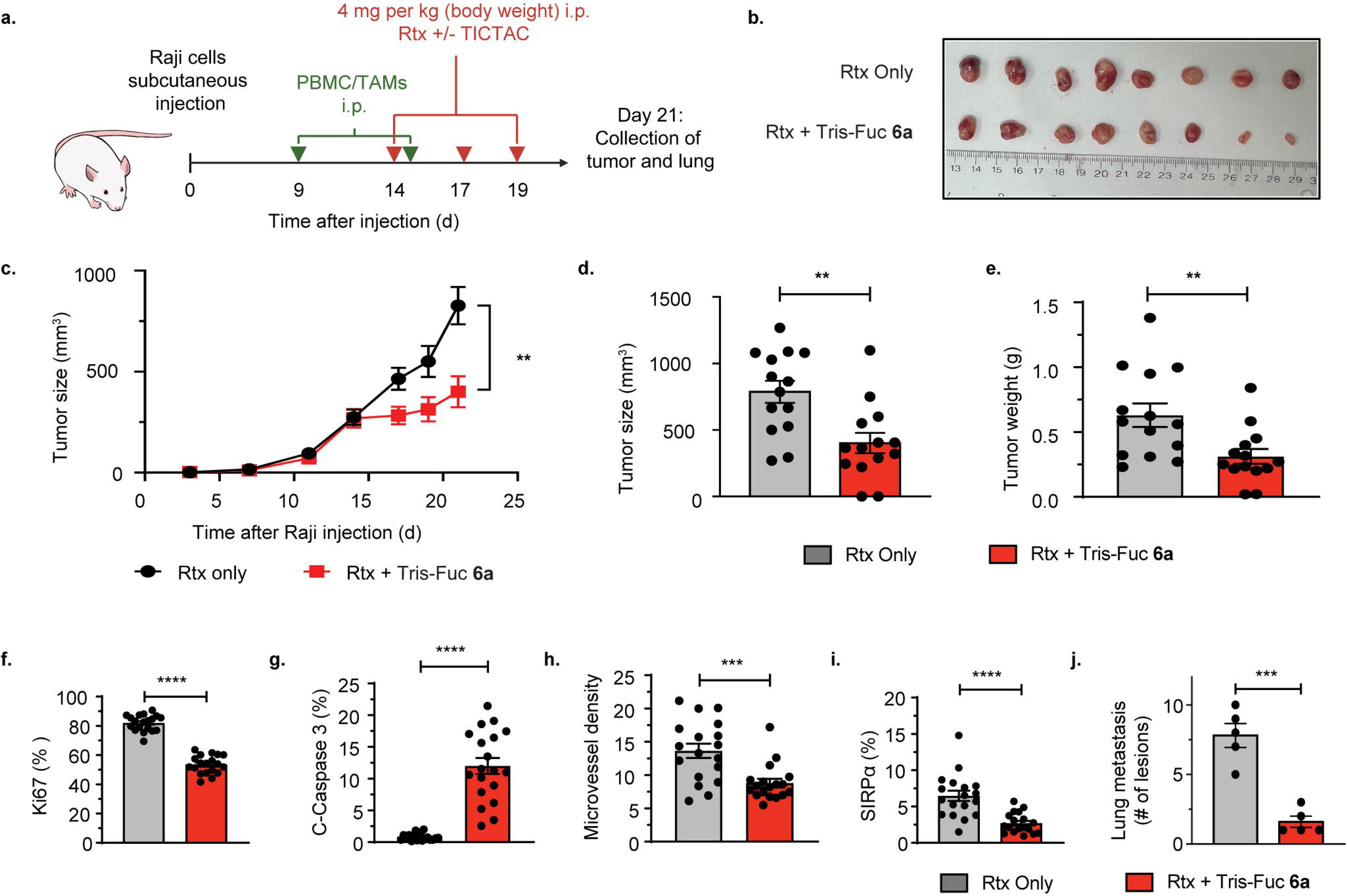
Anti-SIRPα TICTACs reduces tumor burden *in vivo*. **a,** Schematic of the *in vivo* NSG mouse model used to assess the therapeutic efficacy of the anti-SIRPα TICTACs. Raji lymphoma cells were implanted subcutaneously on Day 0. On Day 9 the mice received intraperitoneal (i.p.) injections of human peripheral blood mononuclear cells (PBMCs) supplemented with tumor-associated macrophages. Starting on Day 14, animals were treated every other day with either rituximab (anti-CD20 antibody) or co-treatment of rituximab with anti-SIRPa TICTAC. **b,** Representative images of xenografts comparing the two treatment groups. **c,** Tumor growth curves, **d,** tumor volumes, and **e,** tumor weights. Quantification from three randomly selected high-power fields per tumor: **f,** proliferation index (Ki-67-positive area), **g,** apoptosis index (cleaved caspase-3-positive area), **h,** microvessel density (CD31-positive counts), and **i,** relative SIRPα levels. **j,** Lung metastasis index measured as KU70-positive area per lung section. *P* values were determined by unpaired two-tailed *t*-tests. Statistical significance was defined as *P*<0.05, and the asterisks ** indicates *P*<0.01, *** indicates *P*<0.001, and **** indicates *P*<0.0001.

## Discussion

While numerous strategies have been developed to target tumor cells and immune cells including antibody-drug conjugates and immune checkpoint inhibitors, few methods specifically target *tumor-associated* immune cells. Reprogramming the immune cells within the tumor microenvironment may be advantageous in mitigating irAEs and improving the efficacy and therapeutic window of existing therapies. Toward this objective, we developed tumor immune cell targeting chimeras (TICTACs) that specifically target tumor-associated macrophages via the C-type lectin CD206. While previously reported synthetic ligands for CD206 mostly comprise large mannose-containing glycopolymers, we identified tris-Fuc and tris-3-SO_4_-Gal ligands that potently engage CD206 on their own and in the context of antibody conjugates. Interestingly, we also observed that while the tris-3-SO_4_-Gal ligands bind comparably, if not better, than the tris-Fuc ligands to CD206, the degradation efficacy of the tris-Fuc TICTACs was superior. We hypothesize that the observed discrepancies in potency may arise from structural and functional differences between the sulfated glycan-binding R-type domain and fucose-binding CTLDs of CD206^23,24^. The R-type domain is a single N-terminal unit with a relatively limited surface area for binding. Although this domain efficiently mediates endocytosis of soluble targets, co-endocytosis of membrane-bound targets may require a more precise orientation to achieve effective engagement. In contrast, CD206 contains eight CTLDs, which collectively contribute to higher overall valency. Because these domains rely on multivalent interactions for adequate binding, endocytosis of soluble targets may be slightly less efficient, but co-endocytosis of membrane-bound targets likely benefits from the greater flexibility and reduced orientation constraints afforded by multiple CTLDs. Moreover, the difference in degradation efficacy may also be due to the difference in binding epitope location for the sulfated glycan (membrane-distal) compared to the fucose (membrane-proximal) when engaging CD206, as has been previously observed with membrane-proximal targeting antibody chimeras^52,53^. These TICTAC conjugates were found to induce potent degradation of CD54 and SIRPα from M2-polarized macrophages, with their activity regulated by CD206 expression.

CD206 is a hallmark of alternatively activated macrophages whose expression is potently upregulated by Th2 cytokines (IL4, IL13) and immunosuppressive factors (IL10, CSF1) enriched in the TME^54–56^. While CD206 is also expressed on tissue-resident macrophages, including hepatic Kupffer cells and alveolar macrophages, several lines of evidence support a favorable therapeutic window for TICTACs. First, our data directly demonstrate a large differential in TICTAC activity as a function of CD206 expression: M0 macrophages showed only ∼30% SIRPa degradation compared to >90% in M2-polarized macrophages (**Extended Data Fig. 8a**), establishing that TICTAC activity is gated by CD206 expression level rather than mere receptor presence. Second, TAMs comprise up to 50% of total tumor cell mass in solid tumors^57^, creating a high local density of CD206 targets that strongly favors on tumor activity. Lastly, there is precedent for CD206 targeting as in the case with tilmanocept, which has been safely administered across multiple Phase III trials without significant hepatic or pulmonary toxicities, despite CD206 expression on Kupffer cells and alveolar macrophages^58,59^. Collectively, the expression-dependent activity profile of TICTACs, the abundance of CD206^hi^ TAMs in the tumor microenvironment, and the clinical safety record of CD206-targeted agents support a viable therapeutic window for this modality.

An important consideration for the clinical translation of TICTACs is whether prolonged treatment compromises the native functions of CD206, which plays roles in immune surveillance, pathogen clearance, and collagen homeostasis. Several observations from our study address this concern. Our *in vivo* efficacy study (**Fig. 6**) involved TICTAC dosing every other day for one week (days 14, 17, and 19), constituting repeated *in vivo* exposure without observable adverse effects, providing direct evidence that short-course treatment does not ablate CD206 function. Also, because CD206 constitutively recycles every ∼5 min, with only 10-30% of total receptor present at the cell surface at steady state, this inherently protects against functional depletion, as TICTACs exploit the existing endocytic cycle rather than disrupting it. Moreover, unlike antibody-drug conjugates that deliver cytotoxic payloads, TICTACs do not alter the intrinsic recycling machinery. Consistent with this, we observed preserved surface CD206 levels following TICTAC treatment (**Extended Data Fig. 5**, **Supplementary Fig. S10**) and preserved collagen uptake capacity (**Extended Data Fig. 9 and Supplementary Fig. S15**). Longer-term studies in chronic dosing models will be an important component of future preclinical development to further define the safety profile of this approach.

We also established that TICTACs generated from non-blocking antibodies can still degrade SIRPα in the presence of CD206, which enables antibody selectivity to be decoupled from its blocking function. Moreover, we found that despite leveraging CD206 to mediate uptake, TICTAC treatment does not compromise the innate biological activity or the overall cell surface levels of CD206. The degradative capability of these conjugates is among the highest reported amongst known extracellular degraders, enabling up to 94% degradation of SIRPα in primary human macrophages. A notable feature of TICTACs is the absence of a hook effect across the 0.1-100 nM concentration range tested (**Fig. 3e**). In conventional bifunctional degraders such as PROTACs, the Hook Effect arises because both the target binding warhead and the E3 ligase recruiting moiety have high (nanomolar) affinity^60^. At saturating concentrations, the degrader individually occupies both binding partners, forming two populations of unproductive binary complexes that compete with ternary complex formation. TICTACs have a fundamentally different affinity profile: while the anti SIRPα antibody component binds its target with high nanomolar affinity, the tris fucose ligand binds CD206 with a Kd of ∼4.58 μM by SPR (**Extended Data Fig. 3a**) and an EC50 of 586 nM in a cellular binding assay (**Extended Data Fig. 3b**). The highest TICTAC concentration tested (100 nM) is approximately six-fold below the EC50 for CD206 engagement, meaning that at no point in the tested range does CD206 binding approach saturation. This large affinity asymmetry ensures that the unproductive TICTAC-CD206 binary complexes required to produce a hook effect do not form in appreciable quantities. Moreover, constitutive CD206 recycling continuously refreshes the available receptor pool at the cell surface, further disfavoring binary complex accumulation. Importantly, we demonstrate that TICTACs have a significant effect on the anti-tumor function of M2 macrophages across multiple human donors. While previous targeted protein degradation methods have canonically targeted cancer cells, this work establishes the paradigm of targeting and empowering dysregulated immune cells selectively within the TME.

Additionally, we demonstrated that a single tris-fucose ligand can function as a small molecule CD206 binder even without multivalent display on an antibody. Small molecule binders to CD206 are scarce^61^; previous approaches focused primarily on glycan-conjugated polymers, liposomes, nanoparticles, and other biomaterials. Further, this work presents a rare example of small molecule binders that are not based on mannose yet are highly selective for CD206 in M2-polarized macrophages.

Our strategy for generating a library of antibody conjugates through combinatorial screening of various linker scaffolds and glycan ligands enables the design of a targeting chimera without a pre-optimized small molecule ligand in hand. Additionally, antibody conjugation greatly simplifies the purification process of the small molecule ligands, since size-exclusion techniques such as molecular weight cut-off resins can be used to separate the conjugates from unreacted material. Importantly, these ligands retain their function in the absence of the parent antibody. Thus, we anticipate that this workflow can be utilized to discover novel small molecule binders to other lectins thereby enabling applications in new drug delivery platforms.

Reprogramming the immune system represents a powerful strategy for cancer therapy. As more targeted approaches are being developed for tumor cell-specific therapies, we envision a parallel opportunity to develop tumor immune cell-specific therapies that alleviate irAEs and expand the eligible patient population for cancer immune therapy. TICTACs that leverage CD206 present one such approach for selectively reprogramming TAMs. Further discovery and targeting of immune cell receptors that are upregulated within the TME will expand the scope of this strategy to other immune cell types and checkpoint molecules.

## Acknowledgements

We gratefully acknowledge Dr. Michael Eckert and Jessica Tran (PAN Facility at Stanford) for performing SPR experiments. We also acknowledge Abel Bermudez (Canary Center at Stanford) and Dr. Peter Szijj for assisting with MALDI-TOF-MS characterization. We thank Dr. Jonathan Yang, Chloe Rollock, Cindy Sandoval Espinoza, and Dr. Jessica Stark for experimental assistance and helpful discussions.

## Author Contributions

M.M., D.S.R., and C.R.B. conceived the study. M.M., D.S.R., R.W., E.H., E.E.P., G.M.S., and N.A.T. designed and performed experiments and analyzed data. M.M., C.R.B., and J.D.B. supervised the experimental design, data analysis, and data interpretation. M.M., D.S.R., R.W., and C.R.B. wrote the manuscript. M.M. and C.R.B. provided funding for this study. All authors reviewed and/or revised the manuscript.

## Competing Interests

C.R.B. is a cofounder and Scientific Advisory Board member of Lycia Therapeutics, Palleon Pharmaceuticals, Enable Bioscience, InterVenn Bio, Firefly Bio, Redwood Bioscience (a subsidiary of Catalent), Neuravid Therapeutics, GanNA Bio, TwoStep Therapeutics, and Valora Therapeutics. C.R.B. is a member of the Board of Directors of Eli Lilly and Co., Xaira Therapeutics, and OmniAb. M.M., N.A.T., and C.R.B. are coinventors on a patent application related to this work (U.S. Patent Application No. PCT/US2024/053731). The remaining authors declare no competing interests.

## Funding Statement

C.R.B. discloses support for the research of this work from the National Institutes of Health [grant number GM058867]. M.M. discloses support for the research of this work from the University of Notre Dame [start-up grant]. D.S.R. discloses support from the Connie and Bob Lurie Postdoctoral Fellowship of the Damon Runyon Cancer Foundation [grant number DRG-2526-24]. N.A.T. discloses support from the National Institutes of Health F32 Postdoctoral Fellowship [grant number F32GM143843]. We also acknowledge the National Institutes of Health High End Instrumentation grant [grant number S10OD028697-01] for the Bruker Neo-500 MHz instrument. The remaining authors declare no relevant funding.

## Methods

### General synthetic chemistry procedures

Unless otherwise noted, all reactions were carried out in flame-dried glassware sealed with rubber septa under a nitrogen atmosphere with Teflon-coated magnetic stir bars. Reaction progress was monitored using thin layer chromatography on Millipore Sigma glass-backed TLC plates (250 μm thickness, F-254 indicator) and visualized with 254 nm UV light or stained by submersion in a basic potassium permanganate solution or 5% H_2_SO_4_ in methanol solution. Flash column chromatography was performed on a Biotage instrument using pre-packed silica gel columns. Reagents were purchased in reagent grade from commercial suppliers and used as received, unless otherwise described. Anhydrous solvents (dichloromethane, tetrahydrofuran, methanol, and acetonitrile) were purchased as sure-sealed bottles and used as received, unless otherwise described. See **Supplemental Notes 1** and **2** for detailed synthetic procedures and characterization of compounds.

### TICTAC antibody conjugation

#### General procedure for antibody linker labeling

A 1 mg/mL solution of antibody was buffer exchanged into PBS using a 7 kDa Zeba size-exclusion column. The antibody was reacted with 50 equiv. of NHS-linker (10 mM stock solution in DMSO), and the reaction was incubated overnight at room temperature. The resulting mixture was purified using a 7 kDa Zeba size-exclusion column to yield the conjugate.

#### Example of rabbit IgG-488 conjugation

A 1 mg/mL solution of goat anti-rabbit IgG (Jackson ImmunoResearch, 111-005-144) was buffer exchanged into PBS using a 7 kDa Zeba size-exclusion column. The rabbit IgG antibody was then incubated with 50 equiv. of a 10 mM stock solution (prepared in DMSO) of Alexa Fluor 488 NHS Ester (Thermo Fisher, A20000) overnight at room temperature. After this time, the reaction mixture was passed through twice with a 7 kDa Zeba desalting column (500 µL): first pass which had been equilibrated with Tris-HCl (pH 8.0, 50 mM) to quench any unreacted NHS-linker; second pass which had been equilibrated with PBS.

#### General procedure for Cu-catalyzed click reaction

To a 1 mg/mL solution of the antibody-linker conjugate was added 75 equiv. of the carbohydrate ligand (10 mM stock solution in H_2_O). BTTP-Cu(II) pre-catalyst was generated by mixing CuSO_4_ (aqueous solution) and BTTP (DMSO solution) in a 1:4 molar ratio. 200 μM BTTP-Cu and 4 mM sodium ascorbate (100 mM solution in H_2_O, prepared immediately prior to addition) were added, and the reaction was briefly vortexed. The reaction mixture was allowed to incubate at room temperature for 1 h and was filtered using a 7 kDa or 40 kDa Zeba size-exclusion column.

### Cell culture

All cell lines were purchased from the American Type Culture Collection (ATCC) unless otherwise noted. RAW264.7, J774A.1, BV-2, and U937 were cultured in DMEM supplemented with 10% heat-inactivated fetal bovine serum (HI FBS), 100U/mL Penicillin, and 100 μg/mL Streptomycin. THP-1 cells were cultured in RPMI supplemented with 10% HI FBS, 100U/mL Penicillin, and 100 μg/mL Streptomycin. The CB7BL/6-derived immortalized bone marrow macrophage cell line BMA3.1A7 was a generous gift from Dr. Kenneth Rock (University of Massachusetts Medical School), and the cells were maintained as previously described^62^. Primary murine BMDMs were obtained from Charles River Laboratories and cultured in DMEM supplemented with 10% HI FBS, 100U/mL Penicillin, and 100 μg/mL Streptomycin. Cells were cultured in T-75 flasks (Fisher Scientific) and incubated at 37 °C and 5% CO_2_ and tested negative for mycoplasma quarterly using a PCR-based assay.

### Gene knockout pool

CRISPR-Cas9 mediated knockout cell pool of CD206 in RAW264.7 cells were generated by Synthego Corporation (Redwood City, CA). Cells were electroporated with Streptococcus pyogenes Cas9 (Sp-Cas9) and sgRNAs targeting CD206 using Synthego’s optimized protocol. 48 h post-electroporation, genomic DNA was extracted, PCR amplified and sequenced using Sanger sequencing to verify editing efficacy. Resulting chromatograms were analyzed using Synthego Inference of CRISPR edits software (ice.synthego.com). Mock control cells were generated by electroporating Sp-Cas9 without sgRNA. Cells were cultured in DMEM supplemented with 10% heat-inactivated fetal bovine serum (FBS), 100U/mL Penicillin, and 100 μg/mL Streptomycin.

### Surface plasmon resonance (SPR)

Sensor NTA chip, 350 mM EDTA, 0.5 mM NiCl_2_, N-Hydroxysuccinimide (NHS), 1-Ethyl-3-(3-dimethylaminopropyl) carbodiimide (EDC), and ethanolamine hydrochloride (EA) were obtained from GE Healthcare/Cytiva (Marlborough, MA). SPR experiments were performed on a BIACORE T200 biosensor system (GE Healthcare/Cytiva) at 25 °C. The Sensor Chip NTA (GE Healthcare/Cytiva) was conditioned by injecting 350 mM EDTA regeneration solution over both reference and experimental surfaces for 60 s at a flow rate 30 μL/min. To capture the His-tagged mouse CD206 protein (ACROBiosystems, CD6-H52H9) the experimental NTA flow cell surface was saturated with nickel by injecting nickel solution (0.5 mM NiCl_2_) for 60 s at 10 μL/min, followed by an Extra Wash with 3 mM EDTA to remove any remaining traces of nickel in the system. The His-tagged CD206 protein captured on the Sensor Chip NTA was stabilized by covalent coupling to the chip surface by amine coupling chemistry using *N*-hydroxysuccinimide (NHS) and *N*′-(3-dimethylaminopropyl) carbodiimide hydrochloride (EDC) according to the manufacturer’s instructions. The reference surface was treated the same without the presence of His-tagged CD206 protein. Amino-PEG4-tris-α-L-fucose was diluted in PBS Assay Buffer and injected over the CD206-containing surface for 60 s at different concentrations at a flow rate of 30 μL/min. For each experiment at least 5 different concentrations of analyte molecules were injected over each experimental and control flow cell. Dissociation was allowed to occur at the same flow rate for 150 s followed by Regeneration Buffer (150 mM NaCl/NaAc pH 5) at a flow rate of 50 μl/min for 30 s and Assay Buffer alone at a flow rate of 30 μL/min to allow the baseline to stabilize. All data were corrected for unspecific binding by subtracting the signal measured in the reference flow cell lacking immobilized ligand.

### Soluble target uptake assay via flow cytometry

For uptake experiments, cells were plated (60,000 cells/well in 24-well plates) two days before treatment in complete media supplemented with 25 ng/mL murine IL-4 (Peprotech) for M2-polarization or without any cytokines (M0). Cells were incubated with 250 μL of M2- or M0-media with fluorescent proteins and TICTACs/controls or conjugated/unconjugated dyes for the indicated times (<8 h). For overnight uptake experiments, cells were plated/polarized one day before treatment. After treatment, cells were washed with PBS, then treated with 200 μL of enzyme-free Cell Dissociation Buffer (Gibco) for 10 minutes to dissociate the cells. The resuspended cells were transferred to a pre-chilled 96-well *v*-bottom plate (Corning) and washed with cold blocking buffer (PBS supplemented with 0.5% BSA and 5 mM EDTA) two times, pelleting by centrifugation at 500*g* for 3 min at 4 °C between washes. For experiments requiring co-staining for CD206 levels, Fc-receptors were first neutralized by incubating cells on ice for 5-10 minutes with TruStain FcX (BioLegend) at 2 μg/mL in 100 μL of blocking buffer. The cells were then stained with the fluorescently labeled anti-CD206 antibody in 100 μL of blocking buffer for 30 minutes at 4 °C. Cells were washed two times with blocking buffer, then incubated with Sytox Blue according to the manufacturer’s specifications for 10 min at 4 °C. Flow cytometry was performed on either a MACSQuant Analyzer 10 (Miltenyi Biotec), Novocyte Quanteon (Agilent), or Novocyte Penteon (Agilent), and FlowJo software was used to gate on single cells, live cells, and CD206^hi^/^lo^ cells for analysis. Representative gating strategies are shown in **Supplementary Fig. S16**.

### Cell-surface protein internalization assay via flow cytometry

Cells were plated (60,000 cells/well in 24-well plates) one day before treatment in M2- or M0-media. Cells were incubated with 250 μL of M2- or M0-media with 25 nM TICTACs/controls for the indicated amount of time, washed with PBS, and dissociated with Cell Dissociation Buffer for 5-10 minutes. Resuspended cells were transferred to a 96-well *v*-bottom plate and washed with cold blocking buffer, Fc-neutralized, and incubated with primary antibodies for 30 min at 4 °C. Cells were washed two times with blocking buffer, then incubated with Sytox Blue for 10 min before flow cytometry analysis. Representative gating strategies are shown in **Supplementary Fig. S16**.

### Plasmids and expression of SIRP**LJ** antibodies, LALAPG-119, 136 and 3

LALAPG-119, 136, and 3 plasmid sequences are listed in Supplementary Note 3. Antibodies were expressed in Chinese hamster ovary (CHO) cells, and the purity was >95% by SDS-PAGE.

### Isolation and differentiation of primary human macrophages

LRS chambers were obtained from healthy anonymous blood bank donors (Stanford Blood Center). PBMCs were isolated using SepMate™-50 PBMC isolation tubes (STEMCELL Technologies) and Ficoll-Paque (GE Healthcare Life Sciences) following the manufacturer’s protocols. Monocytes were isolated by plating ∼3 x 10^8^ PBMCs in T-75 flasks with serum-free RPMI and incubating for 1 h, followed by 3x rigorous washes with DPBS (PBS +Ca +Mg) to remove non-adherent cells and ensuring that the final wash results in clear supernatant. The media was replaced with 20 mL IMDM supplemented with 10% Human AB Serum (Gemini), and the cells were allowed to differentiate into macrophages over 7–9 days. To generate M2 macrophages, the media was replaced on day 4 with IMDM supplemented with 10% Human AB Serum, 20 ng/mL IL-4 (Peprotech), 50 ng/mL IL-10, and 50 ng/mL TGF-β. No media changes were conducted for M0 macrophages throughout the differentiation period.

### Western blot analysis

Cells were plated (60,000 cells/well in 24-well plates) one day before treatment in M2- or M0-media. Cells were incubated with 250 μL of M2- or M0-media with 25 nM TICTACs/controls for the indicated amount of time, washed with 3x with PBS, and lysed with RIPA buffer supplemented with protease inhibitor cocktail (Roche), 0.1% benzonase (Millipore-Sigma), and phosphatase inhibitor cocktail (Cell Signaling Technologies) on ice for 30 min. The cells were scraped and transferred to 1.5 mL Eppendorf tubes and centrifuged at 21,000*g* for 15 min at 4 °C. The supernatant was collected, and the concentration was determined using a BSA assay (Pierce). Equal amounts of protein were loaded onto a 4-12% Bis-Tris gel (BioRad) and separated by SDS-PAGE. The gel was transferred to a nitrocellulose membrane, stained with REVERT Total Protein Stain (LI-COR) and blocked with Odyssey Blocking Buffer (PBS) (LI-COR) for 1 h at room temperature. The membrane was then incubated with primary antibody in Blocking Buffer overnight at 4 °C, then washed three times with PBS with 0.1% Tween-20 (PBS-T). The membrane was incubated with secondary antibody for 1 h at room temperature, washed three times with PBST, and visualized with an Odyssey CLx imager (LI-COR). Image Studio (LI-COR) was used to analyze the image and quantify band intensities.

### Confocal microscopy

#### Membrane protein analysis

Cells were plated (20,000 cells per well in an 8-chamber Labtek plate) 3 days prior to treatment. Cells were incubated with 250 μL of complete growth medium with 25 nM TICTACs or control for the indicated amount of time. Cells were then washed 3x with DPBS (300 μL/wash) and fixed with 4% paraformaldehyde in PBS for 10 min at room temperature. Cells were washed 2x with PBS and incubated with primary antibody in 300 μL blocking buffer (PBS + 1% BSA) for 1 h at room temperature. After washing with PBS, cells were incubated with Hoechst stain in PBS for 10 min at room temperature. The wells were aspirated, and the chambers were removed according to the manufacturer’s specifications. 15 μL of mounting medium (Vector Laboratories) was added followed by a coverslip. The slides were cured for 30 min at room temperature and imaged with Nikon A1R confocal microscope using a Plan Fluor x60, 1.30-NA oil objective. The following laser settings were used: 405 nm violet laser, 488 nm blue laser, 561 nm green laser, and 639 nm red laser.

### Phagocytosis assays

On Day 7, M2-polarized human macrophages were washed with PBS and lifted by incubating with 8 mL TrypLE (Thermo Fisher Scientific) for 30 min at 37 °C. The macrophages were pelleted by centrifugation at 400 x *g* for 4 min and resuspended in phenol-red free IMDM supplemented with 10% Human AB serum and M2 cytokine cocktail (20 ng/mL IL-4, 50 ng/mL IL-10, and 50 ng/mL TGF-β). 10,000 macrophages were plated in 100 μL of media in a 96-well flat-bottom plate (Corning) and incubated in a humidified incubator for 1 h at 37 °C. TICTACs or unconjugated antibody were added at 25 nM (in 5 μL volume), and the cells were incubated for 48 h at 37 °C. The wells were aspirated, then replaced with 100 μL IncuCyte medium (phenol-red free RPMI + 10% HI FBS). Raji cells were washed 1x with PBS, then incubated with 1:80,000 diluted pHrodo red succinimidyl ester dye (Thermo Fisher Scientific) in PBS at 37 °C for 30 min on a shaker, washed with 1x PBS, and resuspended in IncuCyte medium. The labeled Raji cells were added to the macrophages (20,000 cells, 90 μL), followed by 10 μL of 20x rituximab stocks (final concentration of 5 μg/mL). Cells were plated by gentle centrifugation (50 x *g,* 1 min) then placed in an IncuCyte S3 Live-Cell Analysis System (Sartorius). Two images per well were acquired at 1 h intervals until maximum signal was reached (at least 4 h). Images were analyzed using a threshold of 1.1, an edge sensitivity of −45, and areas between 100 and 2000 μm^2^. The total red object integrated intensity (RCU x μm^2^/image) was quantified for each well and normalized by dividing by the total confluence (%). This normalized total integrated intensity value is reported for each treatment condition.

### Collagen uptake assay for assessing macrophage function

Collagen polymerization of tissue-culture plates and subsequent macrophage uptake experiments were performed following a previously published protocol^48^. Briefly 10 µg/mL of FITC-collagen (Millipore Sigma, C4361) was partially polymerized on 24-well tissue culture plates for 1 h in the dark. Human donor-derived M2-polarized macrophages (IL4/IL10/TGF-β) were then added to the plates at 100k cells per well. The macrophages were then treated with either: (1) a combination of rabbit IgG-647 antibody (50 nM) and a goat anti-rabbit antibody (with or without conjugation, 25 nM); or (2) a small molecule ligand (Alexa Fluor 647 or tris-Fuc-647 at 25 nM). After allowing the cells to ingest the collagen for 24 h incubation at 37 °C in an incubator, the total collagen uptake and rabbit IgG-647 internalization was then determined by flow cytometry.

### Acquisition parameters for IncuCyte images

IncuCyte S3 Live-Cell Analysis System (Sartorius) was placed within a tissue culture incubator (Thermo Fisher Scientific) maintained at 37 °C with 5% CO_2_. Images were acquired from a 10x objective lens in phase contrast and from a red fluorescence channel (ex. 585 ± 20, em: 665 ± 40, acquisition time: 400 ms). Unless otherwise specified, cells were analyzed by Top-Hat segmentation with 100 μm radius, edge split on, hole fill: 0 μm^2^.

### *In vivo* mouse study

NSG (NOD.Cg-Prkdcscid Il2rgtm1Wjl/SzJ) mice (8-10 weeks, male, n=1, female=13) were obtained from Jackson Laboratory. Mice were housed on a 12-hour light/dark cycle at 20-23 °C and 30-70% humidity. All animal experiments were conducted in accordance with institutional guidelines and were approved by the Stanford University Administrative Panel on Laboratory Animal Care (APLAC). The Raji cells (1×10^6^ cells) were subcutaneously injected into the mice on day 0. PBMCs and TAMs were derived from the same healthy human donors for each experiment. TAMs were generated by isolating monocytes from PBMCs and polarizing them *ex vivo* into M2 macrophages using IL-4 (20 ng/mL), IL-10 (50 ng/mL), and TGF-β (50 ng/mL). PBMCs (10 million/mouse) and TAMs (1:10 TAM:PBMC ratio) were introduced into the mice via intraperitoneal injections weekly beginning on day 9. The control and targeted antibodies (n=7/group, concentration 5 mg/kg) were injected into the mice on day 14 and then injected every other day. At day 21, the mice were sacrificed. Tumor and lung tissues were collected, fixed in formalin, and embedded in paraffin for downstream analysis.

### Hematoxylin and Eosin (H&E) staining

Tissue sections were processed for H&E staining following as previously described^63,64^. After baking at 60 °C for 30 min, slides were sequentially incubated twice for 5 minutes in each of the following solutions: Clearify™ (StatLab, SKU: CACLELT), 100% ethanol, 95% ethanol, and 75% ethanol. Sections were stained with Ehrlich’s hematoxylin for 2 minutes. Following a 3-minute incubation in sodium bicarbonate (50 mM) solution, slides were counterstained with eosin for 30 seconds and rinsed under tap water for three times. Dehydration was performed by immersing slides twice for 2 minutes each in 95% ethanol, 100% ethanol, and xylene. Slides were then mounted with coverslips.

### Immunohistochemistry (IHC) staining

IHC staining was carried out using antibodies against CD31, SIRPα, Ki67, and cleaved caspase-3 as previously described^63,64^. Tissue sections were first baked at 60 °C for 30 min, then deparaffinized and rehydrated through two 5-minute incubations in Clearify™, followed by two 5-minute washes in 100% ethanol and 95% ethanol, and a single 5-minute wash in 70% ethanol. Antigen retrieval was performed by boiling slides in 0.01 M citrate buffer (pH 6.0) for 30 minutes. After cooling, sections were blocked in PBS containing 5% goat serum and 5% BSA for 1 hour at room temperature. Primary antibodies were diluted in PBS with 0.5% BSA and applied overnight at 4 °C. The following day, slides were washed three times in TBST and incubated with 0.3% hydrogen peroxide for 15 min to block endogenous peroxidase activity. A secondary antibody was applied in PBS with 0.05% BSA, followed by detection using DAB substrate (Vector Laboratories, Cat. No. SK-4100). The anti-CD31 antibody was chosen for its specificity for mouse CD31, and the anti-SIRPα antibody was chosen for its specificity for human SIRPα. Slides were counterstained with hematoxylin for 5 minutes and incubated in sodium bicarbonate (50 mM) solution for 3 minutes. Dehydration was carried out by immersing the slides twice for 5 minutes each in 75%, 95%, and 100% ethanol, followed by xylene. Finally, slides were mounted with coverslips.

### Proteomics sample preparation

Macrophage samples cultured on 24-well plates were washed three times with cold (4 °C) PBS (pH 7.4) to remove cell culture media. Following PBS washes, the macrophage samples were lysed in RIPA buffer containing 25 mM Tris-HCl (pH 7.6), 150 mM NaCl, 1% (m/V) NP-40, 1% (m/V) sodium deoxycholate, 0.1% (m/V) and sodium dodecyl sulfate (SDS), and 1x Halt protease inhibitor cocktail (78430, Thermo Fisher Scientific). The cell lysates were then processed using a suspension trapping (S-Trap) microcolumn (Protifi, C02-micro-80, ≤ 100 μg) following vendor protocols as described by Protifi (Fairport, NY, USA) with slight modifications. Briefly, protein samples were first reduced using 20 mM tris(2-carboxyethyl)phosphine (TCEP) at 37°C for 15 min and then alkylated using 40 mM 2-chloroacetamide for 20 min at room temperature. The denatured, non-digested proteins were then acidified using phosphoric acid (85 % w/v) to a final phosphoric acid concentration of 2.5 % w/v before being loaded on the S-trap microcolumn. The bound proteins were washed by repeated centrifugation using 100 mM triethylammonium bicarbonate (TEAB) in 90% methanol to remove MS incompatible buffer salts and detergents. The protein samples were then digested on the S-trap using 50 µL of a 1:20 enzyme to substrate mass ratio of Trypsin (Promega, V5280) at 37 °C overnight for 16 h. The resulting peptides were then eluted from the S-trap using 40 µL of 50 mM TEAB, followed by 40 µL of 0.2% formic acid (FA), and finally with 40 µL of 50% acetonitrile (ACN). The pooled elution mixtures were then dried using a Labconco CentriVap vacuum concentrator (Kansas City, MO, USA). Dried peptides were resuspended in 12 μL of 0.2% FA, 2% ACN and peptide concentrations were measured by absorbance at 205 nm using a Nanodrop (Thermo Fisher Scientific) prior to LC/MS analysis.

### LC-MS/MS analysis

Reconstituted peptides were injected and separated using a trap-and-elute nanoflow method over an Aurora Ultimate TS 25 cm nanoflow UHPLC column (25 cm x 75 μm ID, 1.7 μm C18; IonOpticks, AUR3-25075C18-TS) analytical column connected to a Dionex Ultimate 3000 RPLC nano system (Thermo Fisher Scientific) with an integrated loading pump using mobile phases A (water with 0.2% FA) and B (ACN with 0.2% FA). Peptides were loaded onto a trap column (Acclaim PepMap 100 C18, 5 μm particles, 20 mm length, Thermo Fisher Scientific, 164213) at a flow rate of 5 µL/min under 100 % A which was then put in line with the analytical column 5.5 min into the acquisition. The peptides were then separated at 300 nL/min using the following elution gradient: 2 % B for 6 min, 2 % to 5 % B from 6 to 6.5 min, 5 % to 28 % B from 6.5 to 66.5 min, 28 % to 90 % B from 66.5 to 71 min, isocratic flow at 90 % B from 71 to 75 min, and a re-equilibration at 2 % B for 15 min for a total analysis time of 90 min. Eluted peptides were analyzed on an Orbitrap Fusion Tribrid MS system or an Orbitrap Ascend Structural Biology Tribrid MS system (Thermo Fisher Scientific). Precursors were ionized using an EASY-Spray ionization source (Thermo Fisher Scientific) set at 1.85 kV compared to ground, and the column was held at 45 °C. The inlet capillary temperature was set at 325 °C. Data-dependent acquisition (DDA) included MS1 scans acquired over a mass range of 350-1350 *m/z*, with a resolution of 60,000 at 200 *m/z*, normalized automatic gain control (AGC) target of 250% with absolute AGC value of 1 E6, a maximum injection time (IT) of 100 ms, and RF lens at 60%. DDA MS2 scans were then collected with an isolation window of 2 m/z, a normalized AGC target of 200%, with absolute AGC value of 1 E5, a maximum IT of 50 ms, Orbitrap resolution of 15,000 at 200 *m/z*, normalized HCD collision energy of 30%, and a scan range of set to first mass at 120 *m/z*. Monoisotopic precursor selection was enabled for peptide isotopic distributions, precursors of *z* = 2 to 5 were selected for data-dependent MS/MS scans for 2 s of cycle time, and dynamic exclusion was set to 30 s with a ± 10 ppm window set around the precursor monoisotopic ion.

### LC-MS/MS data analysis

Raw data were processed using FragPipe^65^ (v22.0) with MSFragger v4.1, Philosopher v5.1, and IonQuant v1.10.27 for FDR-controlled match between runs (MBR). MSFragger default parameters were used for the “LFQ-MBR” workflow. Peptide spectral matches were made against a target-decoy human reference proteome (UP000005640) database downloaded from Uniprot (accessed 23 July 2023). Peptides and proteins were filtered to a 1% FDR. Further post-processing and figure generation were carried out in R using the FragPipeAnalystR package^66^. For quantitative comparisons, protein intensity values were log_2_-transformed before further analysis, and missing values were imputed from a normal distribution using the Perseus method^67^ and differential expression analysis was performed using Limma^68^. All *P*-values and adjusted *P*-values are computed using the Benjamini–Hochberg method^69^.

### Matrix-assisted laser desorption/ionization-mass spectrometry

A 1 mg/mL solution of the antibody conjugate was buffer exchanged into ddH_2_O using a 7K Zeba size-exclusion column to remove excess salts. Samples were prepared by mixing 2 μL of sinapinic acid (SPA) matrix (10 mg/mL in 0.1% trifluoroacetic acid and 50% acetonitrile) and 2 μL of the antibody sample. The mixture was vortexed, and 1 μL was loaded onto a MALDI stainless steel plate. The sample was dried at room temperature for 15 min, and the MALDI-MS was acquired by AB SCIEX TOF/TOF and a 5800 CovalX High Mass Detector with a mass range of 10,000 – 250,000 Da and a fixed laser intensity of 5,900. Greater than 3 scans were taken per sample, and the spectra was analyzed, averaged, and plotted using MALDIquant^70^.

### Statistical analysis and software

Statistical analysis was performed in GraphPad Prism (version 9). For SPR data, association then dissociation models were used to calculate the dissociation constant. For EC_50_ data, one-site total and nonspecific binding models were used to determine the apparent dissociation constant for specific binding. Flow cytometry data was analyzed using FlowJo software (version 10.8.1). Confocal microscopy images were processed using ImageJ 1.53t. NMR data was analyzed using MestreNova software (version 14.2.3). Figures were created using BioRender.com and Adobe Illustrator software (version 30.1).

## Data Availability Statement

The data that support the findings of this study are generated and analyzed within the published article and its Supplementary Information files. Proteomics data have been deposited to the MassIVE repository (MSV000101189) and the ProteomeXchange Consortium via the PRIDE partner repository with the dataset identifier PXD075936. Uncropped, unprocessed images for all Western blots and gels are provided as Source Data with this paper. Any additional source data are available from the corresponding author upon reasonable request.

## Notes

### Summary of Updates

Text edits have been made to expand the discussion, and revisions were made to the figures. Author affiliations have also been updated.

