## Supplementary Information for "Tumor Immune Cell Targeting Chimeras (TICTACs) reprogram tumor-associated macrophages"

### Table of Contents

Supplementary Table 1 ……………………………………………………………………..…….2
Supplementary Table 2 ……………………………………………………………………..…….3
Supplementary Figures 1-16 ……………………………………………………………………4-19
Unprocessed Gels and Western blot images ………………………………………..….20-22
Supplementary Note 1: General methods and instrumentation …………………………….23-24
Supplementary Note 2: Synthesis and characterization of compounds ……………..…….25-27
$^{1}H$ and $^{13}C$ NMR spectra ……………………………………………………………………..28-29
Supplementary Note 3: Amino acid and DNA sequences for LALAPG-119, 136, and 3 ..30-33
Supplementary References ……………………………………………………………………..34

### Supplementary Table 1. Antibody Information and Concentrations

| Antibody | Source (Cat #) | Usage and Dilution |
|---|---|---|
| Hamster anti-CD54 (3E2) | BD-Biosciences (550287) | Functional |
| Rat anti-CD54 | BioXCell (BE0020-1) | Flow cytometry (3 µg/mL) |
| Rabbit anti-ICAM1 [EPR16608] | Abcam (ab179707) | WB, 1:1000 |
| Rabbit anti-SIRPα [EPR16264] | Abcam (ab191419) | WB, 1:1000 |
| Anti-beta tubulin antibody loading control | Abcam (ab6046) | WB, 1:1000 |
| Rat anti-mouse IL-4 | BioXCell (BE0199) | Functional |
| Goat anti-rabbit IgG | Jackson ImmunoResearch (111-005-144) | Functional |
| Polyclonal rabbit IgG | BioXCell (BE0095) | Functional |
| Rat anti-CD206 APC | Thermo Fisher (17-2061-82) | Flow cytometry (2 µg/mL) |
| Mouse anti-CD206 APC | Thermo Fisher (17-2069-42) | Flow cytometry (5 µL/test) |
| Rat anti-CD206 Alexa Fluor 488 | Biolegend (141710) | Flow cytometry (5 µg/mL) |
| Rabbit anti-CD206 | Cell Signaling Technology (24595) | WB, 1:1000 |
| Mouse anti-CD172g PE | Biolegend (336606) | Flow cytometry (5 µg/mL) |
| Mouse anti-CD3 Pacific Blue (OKT3) | Biolegend (317314) | Flow cytometry (5 µg/mL) |
| InVivoSIM anti-human CD20 (Rituximab Biosimilar) | BioXCell (SIM0008) | Phagocytosis (5 µg/mL) |
| Mouse anti-CD45 | BioXCell (BE0300) | Flow cytometry (2 µg/mL) |
| Mouse anti-CD71 | BioXCell (BE0331) | Flow cytometry (2 µg/mL) |
| Rat anti-CD80 | BioXCell (BE0365) | Flow cytometry (2 µg/mL) |
| Rat anti-TIM-3 | BioXCell (BE0115) | Flow cytometry (2 µg/mL) |
| Rat anti-CSF1R | BioXCell (BE0213) | Flow cytometry (2 µg/mL) |
| TruStain FcX (anti-mouse CD16/32) | Biolegend (101320) | Flow cytometry (2 µg/mL) |
| Rat IgG2a, κ Isotype Ctrl | Biolegend (400501) | Flow cytometry (2 µg/mL) |
| Human BD Fc Block | BD Biosciences (564219) | Flow cytometry (5 µg/mL) |
| IRDye 800 CW Goat-anti-rabbit IgG (H+L) | LI-COR (926-32211) | WB, 1:10000 |

### Supplementary Table 2. Information for Chemical Reagents and Stains

| Reagent | Source (Cat #) |
|---|---|
| 2-Azidoethyl α-L-fucopyranoside | Synthose Inc. (AF750) |
| Propargyl α-L-fucopyranoside | Synthose (PF993) |
| 2-Azidoethyl α-D-mannopyranoside | Synthose Inc. (AM482) |
| α-D-mannopyranosyl azide | Synthose Inc. (MM947) |
| 3-[2-[2-(Aminooxy)acetyl]hydrazinocarbonyl]propyl 2α-mannobioside | Synthose Inc. (AM289) |
| 2-Azidoethyl 3,6-di-O-(α-D-mannopyranosyl)-α-D-mannopyranoside | Synthose Inc. (AM229) |
| 3 Fucosyllactose-Nac-propargyl | Elicityl (GLY060-NPR) |
| β-D-galactose pentaacetate | Sigma Aldrich (134031) |
| Alexa Fluor 488 NHS Ester | Thermo Fisher (A20000) |
| Alexa Fluor 647 NHS Ester | Thermo Fisher (A20006) |
| APDye Fluor 647 acid | AxisPharm (AP15222) |
| Azidoacetic acid NHS ester | BroadPharm (BP-22467) |
| Azido-PEG4-NHS ester | BroadPharm (BP-20518) |
| Azido-PEG12-NHS ester | BroadPharm (BP-22855) |
| Propargyl-PEG4-NHS ester | BroadPharm (BP-21612) |
| Ald-Ph-PEG4-NHS ester | BroadPharm (BP-20558) |
| Endo-BCN-PEG4-NHS ester | BroadPharm (BP-24010) |
| NHS-PEG5-tris-alkyne | Conju-Probe, LLC (CP-2209) |
| Amino-PEG4-tris-alkyne | Conju-Probe, LLC (CP-2210) |
| NHS-PEG5-tris-PEG3-azide | Conju-Probe, LLC (CP-2214) |
| NHS-PEG5-tris-PEG3-DBCO | Conju-Probe, LLC (CP-2235) |
| Sytox™ Blue Dead Cell stain, for flow cytometry | Thermo Fisher (S34857), 1:2000 |
| Sytox™ Red Dead Cell stain, for flow cytometry | Thermo Fisher (S34859), 1:2000 |
| pHrodo™ Red, succinimidyl ester | Thermo Fisher (P36600) |
| Revert™ 700 total protein stain for Western blot normalization | LICOR (926-11011) |
| FITC-Conjugated Bovine Collagen Type I | Millipore Sigma (C4361) |

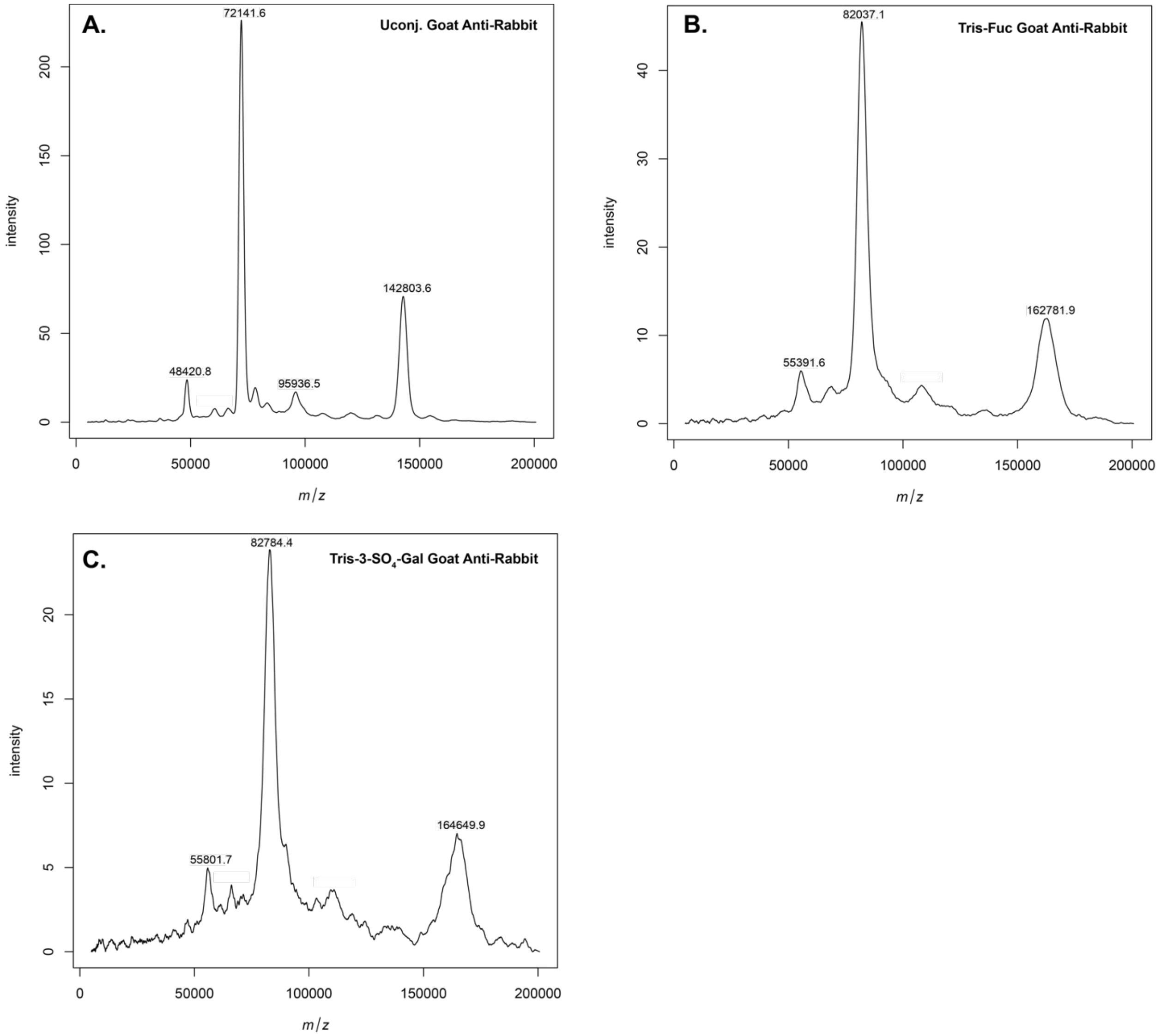


**Supplementary Figure 1.** MALDI-MS characterization of goat-anti-rabbit antibody conjugates. The average ligand antibody ratio (LAR) was determined by subtracting the intact mass of the conjugated antibody from the unconjugated antibody and dividing by the molecular weight of the ligand: tris fucose = 1237.57, tris-3-$SO_4$-Gal = 2201.49. **A,** Combined spectrum of unconjugated goat-anti-rabbit antibody; **B,** Combined spectrum of tris-fucose conjugated goat-anti-rabbit antibody **1a**, where the average LAR is 16; **C,** Combined spectrum of tris-3-$SO_4$-Gal conjugated goat anti-rabbit antibody **1b**, where the average LAR is 10.

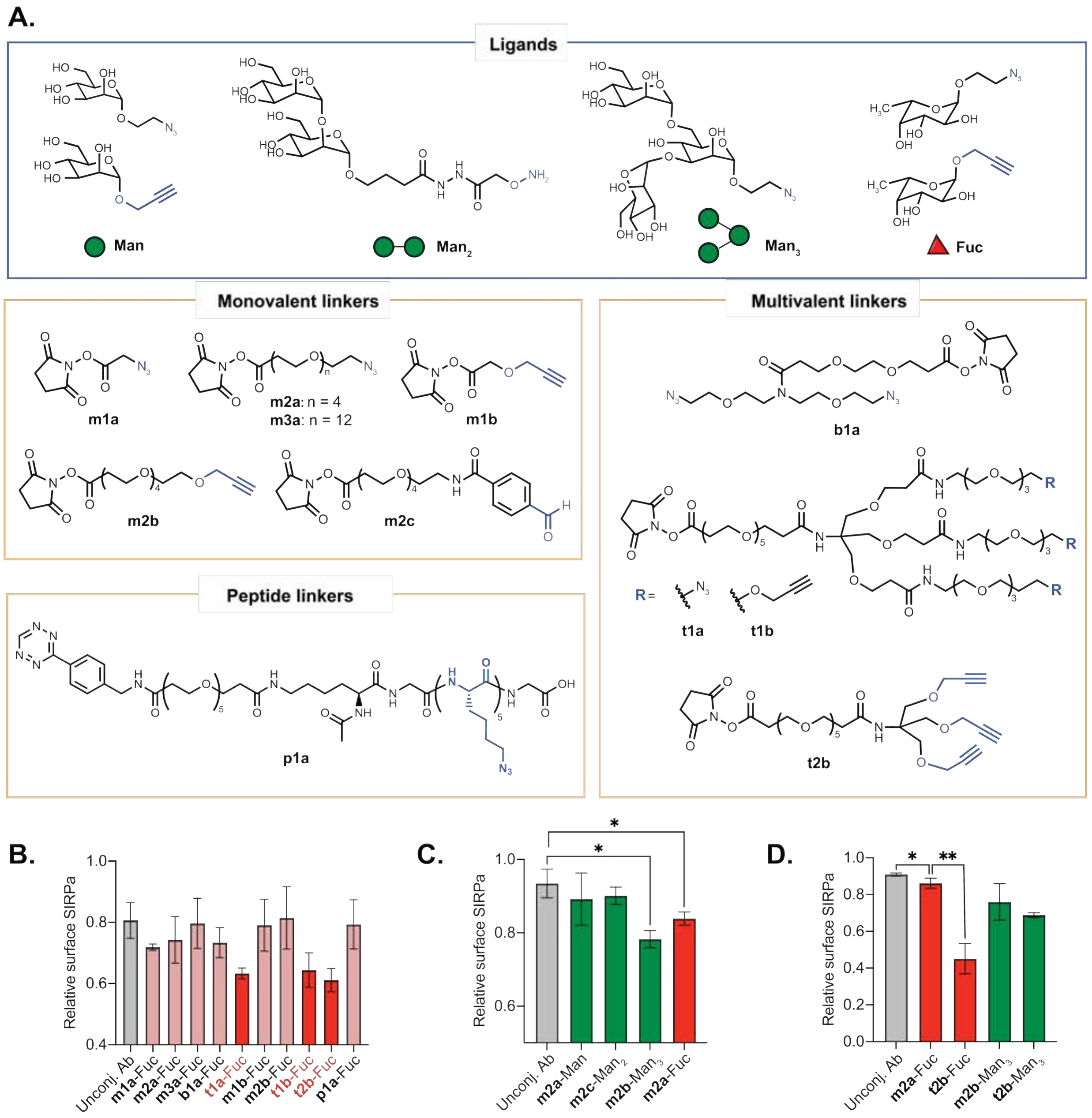


**Supplementary Figure S2. A,** Subset of linkers that were examined in head-to-head comparisons of Fuc and Man ligands. **B,** Comparison of monovalent, bivalent, or pentavalent scaffolds for the Fuc ligand conjugated to an anti-SIRPα antibody (P84 clone). M2-polarized RAW264.7 cells were treated with 25 nM of each conjugate and incubated for 48 h. From this screen, it was found that the trivalent dendron scaffolds (t1a, t1b, and t2b) consistently outperformed either the monovalent, bivalent, or pentavalent scaffolds. **C,** Screen comparing various Man ligands (Man, Man2, Man3) to Fuc using monovalent linkers. All were conjugated to an anti-SIRPα antibody (P84 clone). M2-polarized RAW264.7 cells were treated with 25 nM of each conjugate and incubated for 24 h. Fuc and Man3 were found to be the optimal ligands. **D,** Screen comparing the effect of valency (monovalent vs trivalent) on degradative ability using Man and Fuc ligands downselected from B and conjugated to an anti-SIRPα antibody (P84 clone). The trivalent Tris-Fuc ligand showed superior degradation compared to tris-Man.

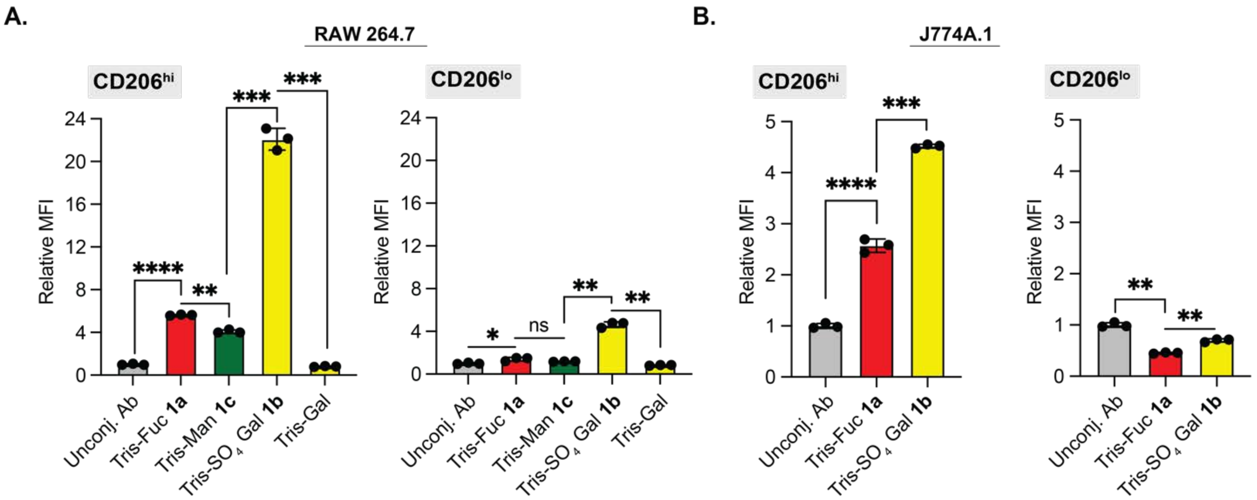


**Supplementary Figure S3.** Additional data for TICTAC-mediated internalization of fluorescent rabbit IgG into M2-polarized macrophages. **A,** Extended time periods (8 h) improves TICTAC-mediated uptake into RAW264.7 cells. **B,** Tris-fucose and Tris-3-$SO_4$-Gal TICTACs induce uptake of fluorescent IgG in M2-polarized J774A.1 cells. Error bars represent the SD from 3 independent experiments. *P* values were determined by Welch's two-tailed *t*-tests. Statistical significance was defined as $P<0.05$, and the asterisks * indicates $P<0.05$, ** indicates $P<0.01$, *** indicates $P<0.001$, and **** indicates $P<0.0001$.

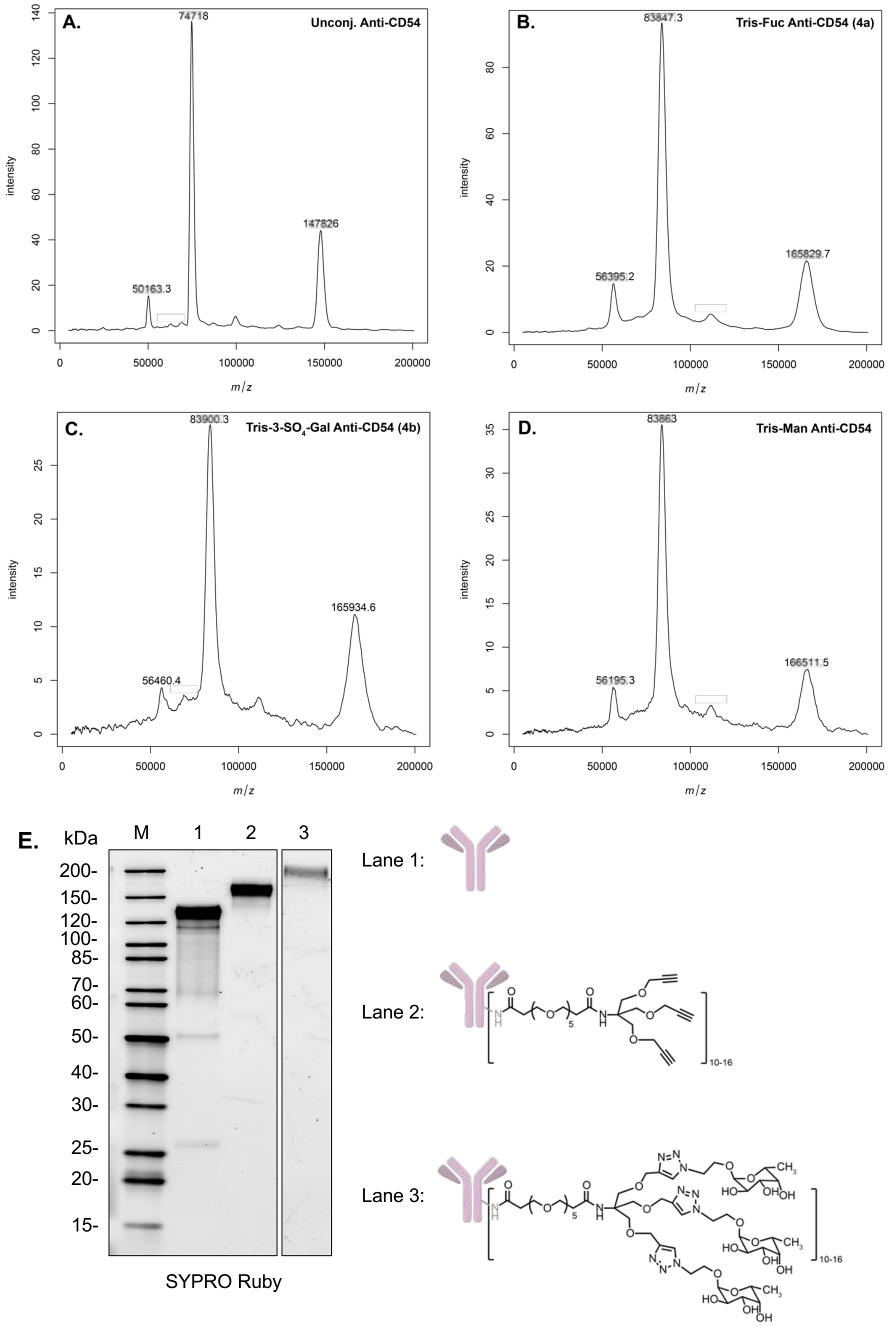


**Supplementary Figure S4.** MALDI-MS characterization of anti-CD54 antibody conjugates. The average ligand antibody ratio (LAR) was determined by subtracting the intact mass of the conjugated antibody from the unconjugated antibody and dividing by the molecular weight of the ligand: tris fucose = 1237.57, tris-3-$SO_4$-Gal = 2201.49, tris mannose = 1285.57. **A,** Combined spectrum of unconjugated anti-CD54 antibody; **B,** Combined spectrum of tris-fucose conjugated anti-CD54 antibody **4a**, where the average LAR is 15; **C,** Combined spectrum of tris-3-$SO_4$-Gal anti-CD54 antibody **4b**, where the average LAR is 9; **D,** Combined spectrum of tris-mannose anti-CD54 antibody, where the average LAR is 15. **E,** SDS-PAGE analysis of unconjugated (lane 1), NHS-PEG5-tris-alkyne conjugated (lane 2), and tris-fucose conjugated (lane 3) anti-CD54 antibody. Total protein visualized by SYPRO Ruby stain. M, molecular weight marker.

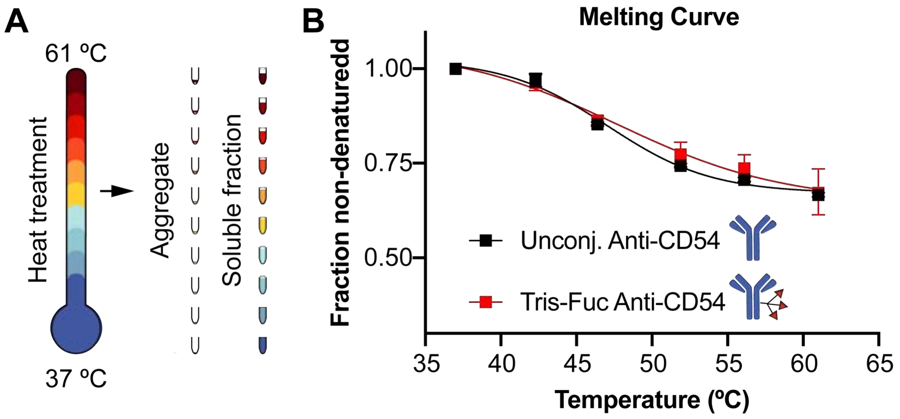


**Supplementary Figure S5**. Thermal proteome profiling (TPP) analysis of Anti-CD54 TICTAC performed according to a previously published method[1]. **A,** Unconjugated Anti-CD54 antibody or Anti-CD54 Tris-Fuc TICTACs were heated in parallel for 3 minutes to 37, 42, 46.4, 51.9, 56.1, and 61 °C, followed by a 3-minute incubation time at room temperature. Insoluble and aggregated proteins were removed by filtration using a multiscreenHTS-HV 0.45-µm 96-well filter plate with PVDF membrane (Merck Millipore). **B,** The flow-through was collected and measured to generate the melting curves. No significant change in melting behavior is observed between the unconjugated antibody (black squares) and the Tris-Fuc TICTAC (red squares) suggesting that Tris-Fuc conjugation does not significantly promote antibody aggregation. Error bars represent the SD from 3 independent experiments.

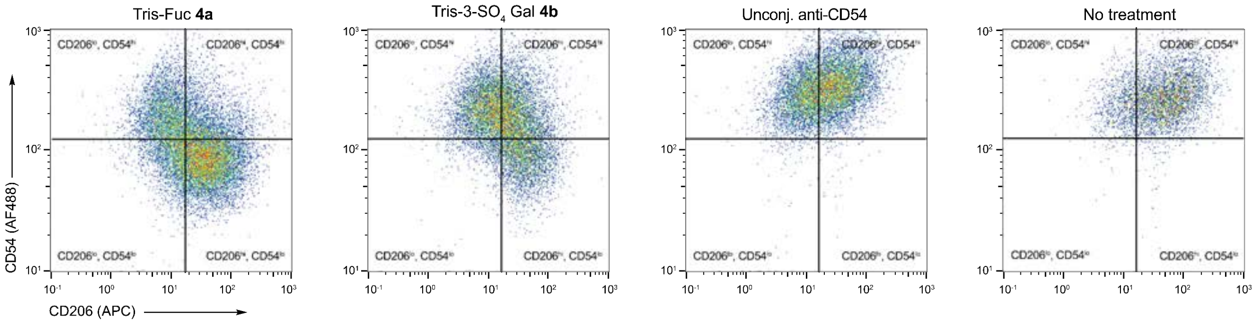


**Supplementary Figure S6.** Correlation between CD206 expression levels and CD54 after treatment of M2-polarized RAW264.7 cells with 25 nM anti-CD54 **4a**, **4b**, or unconjugated antibody for 24 h, measured by live cell flow cytometry.

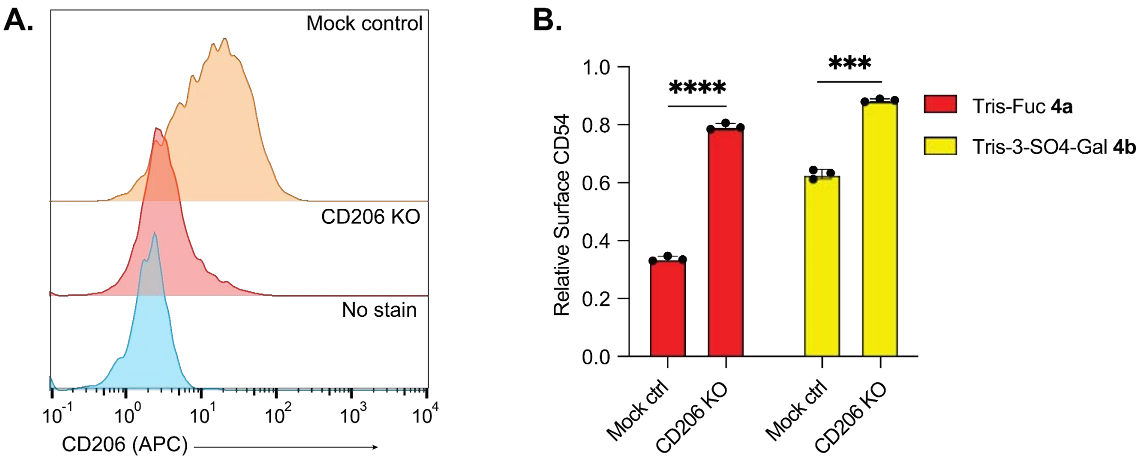


**Supplementary Figure S7.** CD206 knockout greatly attenuates activity of anti-CD54 TICTACs. **A,** Mock control and CD206 knockout RAW264.7 cells were polarized with 25 ng/mL IL-4 for 48 h, then stained for CD206 (APC). **B,** Treatment of CD206 knockout RAW264.7 with **4a** or **4b** over 24 h results in significantly attenuated CD54 downmodulation compared to the mock control. Reported values are normalized to the unconjugated anti-CD54 control. Error bars represent the SD from 3 independent experiments, and *P* values were determined by Welch's two-tailed *t*-tests. Statistical significance was defined as *P*<0.05, and the asterisks * indicates *P*<0.05, ** indicates *P*<0.01, *** indicates *P*<0.001, and **** indicates *P*<0.0001.

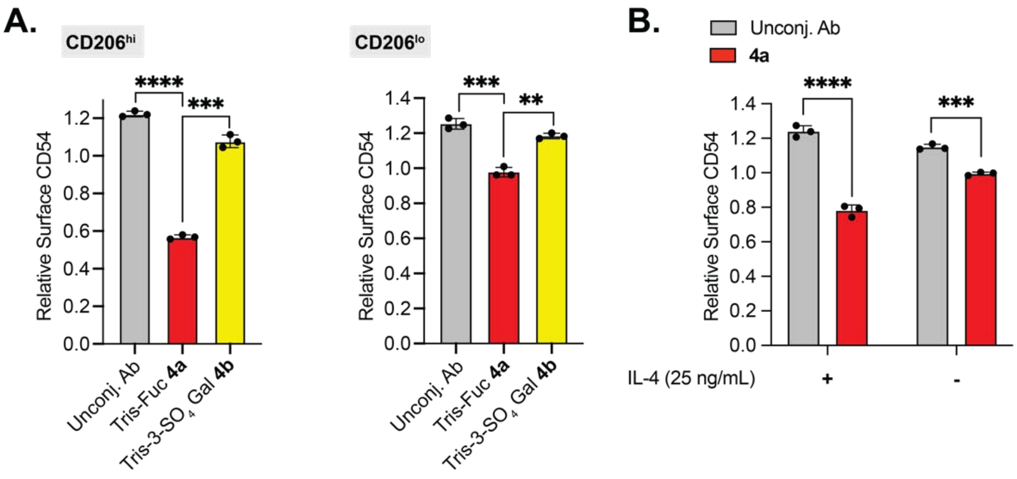


**Supplementary Figure S8.** Anti-CD54 TICTACs downmodulate surface CD54 in J774A.1 cells. **A,** Depletion of cell-surface CD54 in M2-polarized J774A.1 cells determined by live cell flow cytometry following 24 h of treatment with 25 nM Tris-Fuc **4a**, Tris-3-SO4-Gal **4b**, or the unconjugated anti-CD54 antibody. **B,** Depletion of cell-surface CD54 in J774A.1 cells that have been polarized with IL-4 (25 ng/mL) or not following treatment with 25 nM unconjugated antibody or **4a**. Error bars represent the SD from 3 independent experiments, and $P$ values were determined by Welch's two-tailed $t$-tests. Statistical significance was defined as $P<0.05$, and the asterisks * indicates $P<0.05$, ** indicates $P<0.01$, *** indicates $P<0.001$, and **** indicates $P<0.0001$.

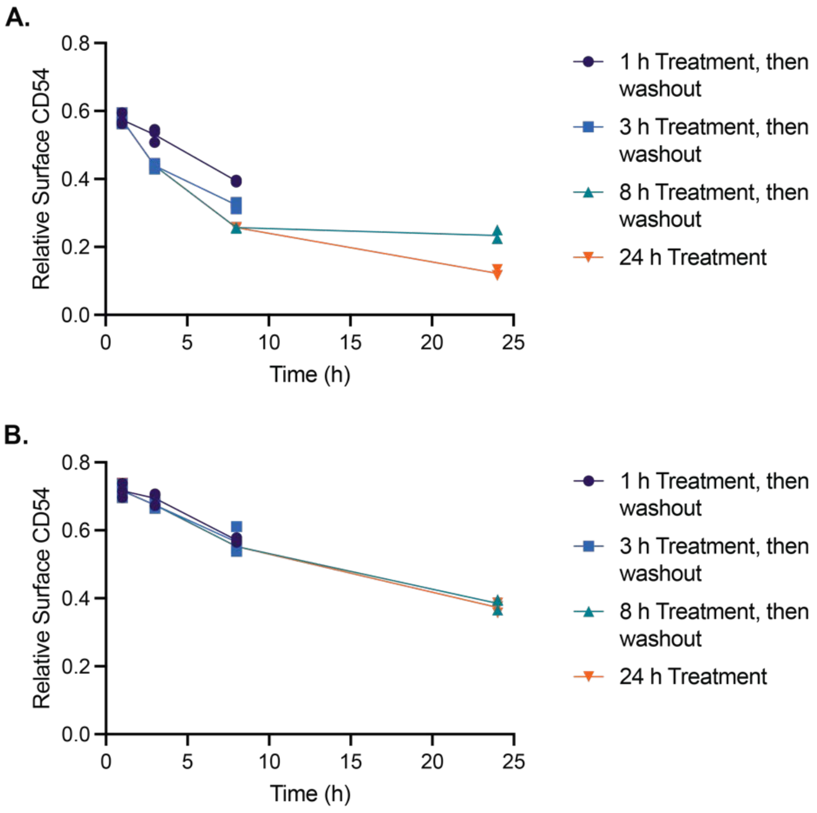


**Supplementary Figure S9.** Washout experiment with M2-polarized RAW264.7 cells where cells were first exposed to media containing **4a**, **4b**, or the unconjugated antibody for 1 h, 3 h, or 8 h (treatment time), then allowed to grow in TICTAC or antibody-free media for varying time periods (washout period). Cell-surface CD54 levels were measured by live-cell flow cytometry. **A,** Washout experiments for **4a**, where surface CD54 levels are normalized to the unconjugated antibody control. **B,** Washout experiments for **4b**, where surface CD54 levels are normalized to the unconjugated antibody control.

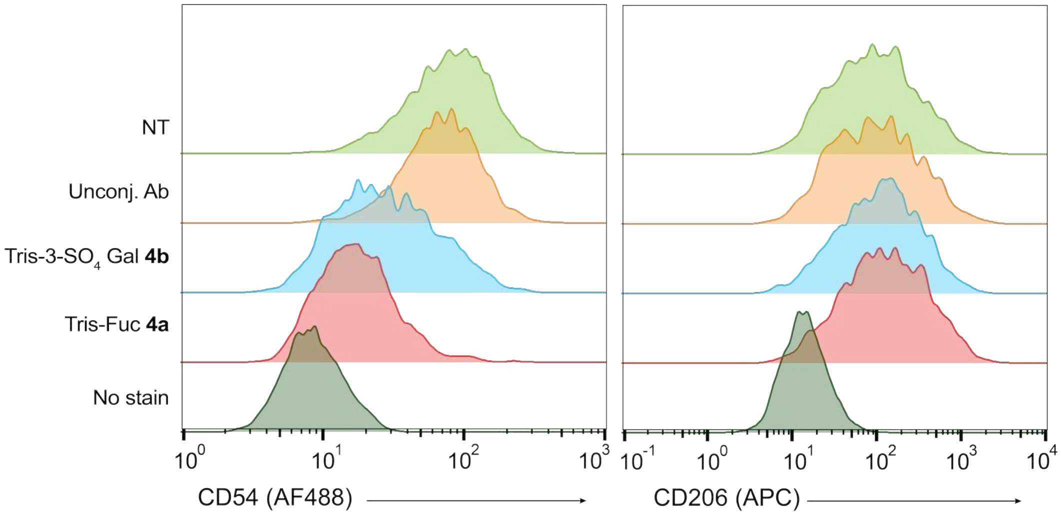


**Supplementary Figure S10.** C57BL/6 BMDMs were treated with anti-CD54 (clone 3E2) TICTACs (tris-3-SO4 Gal 4b or tris-Fuc **4a**) or unconjugated control for 24 h. Cell surface CD54 and CD206 were simultaneously measured after the experiment using flow cytometry, which showed that TICTAC-mediated CD54 degradation does not affect cell surface CD206 levels. Western blot data from this experiment is shown in Fig. 3K.

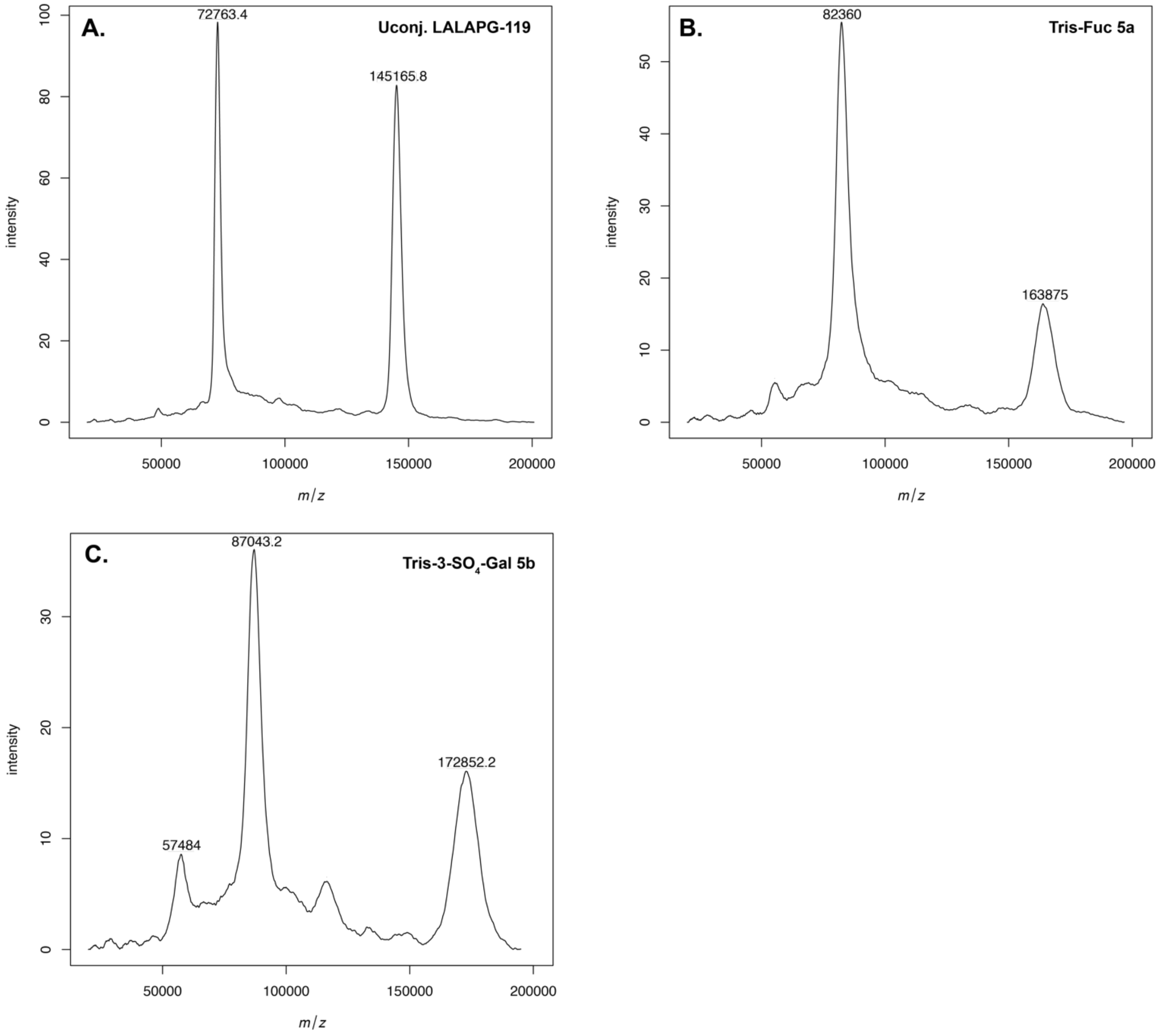


**Supplementary Figure S11.** MALDI-MS characterization of anti-SIRPα antibody conjugates. **A,** Combined spectrum of unconjugated LALAPG-119; **B,** Combined spectrum of tris-fucose conjugated anti-SIRPα antibody 119 (**5a**), where the average LAR is 15; **C.** Combined spectrum of tris-3-$SO_4$-Gal anti-SIRPα antibody 119 (**5b**), where the average LAR is 13.

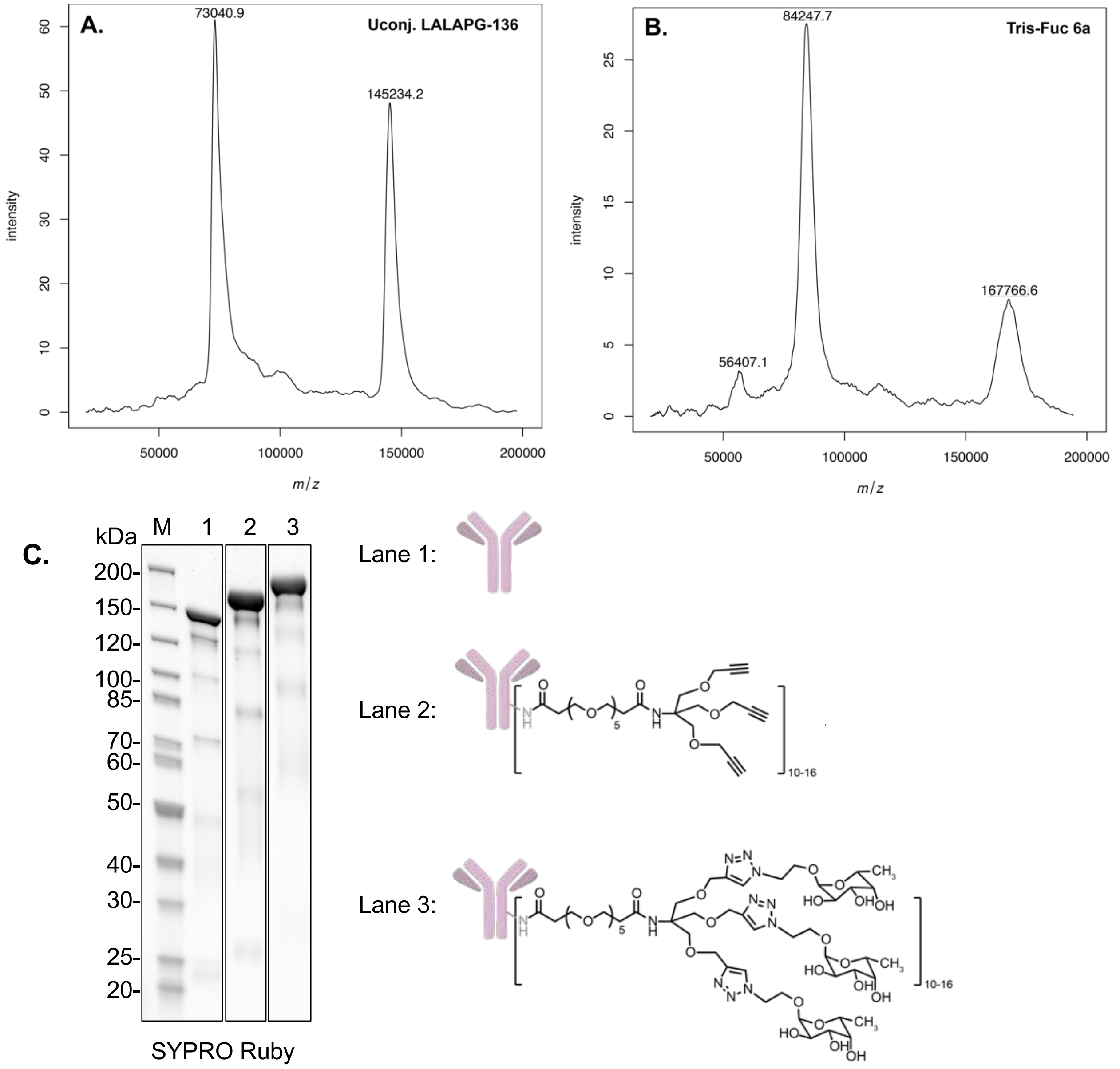


**Supplementary Figure S12.** MALDI-MS characterization of anti-SIRPα antibody conjugates. **A,** Combined spectrum of unconjugated LALAPG-136; **B,** Combined spectrum of tris-fucose conjugated anti-SIRPα antibody 136 (**6a**), where the average LAR is 18. **C,** SDS-PAGE analysis of unconjugated (lane 1), NHS-PEG5-tris-alkyne conjugated (lane 2), and tris-fucose **6a** conjugated (lane 3) anti-SIRPα antibody. Total protein visualized by SYPRO Ruby stain. M, molecular weight marker.

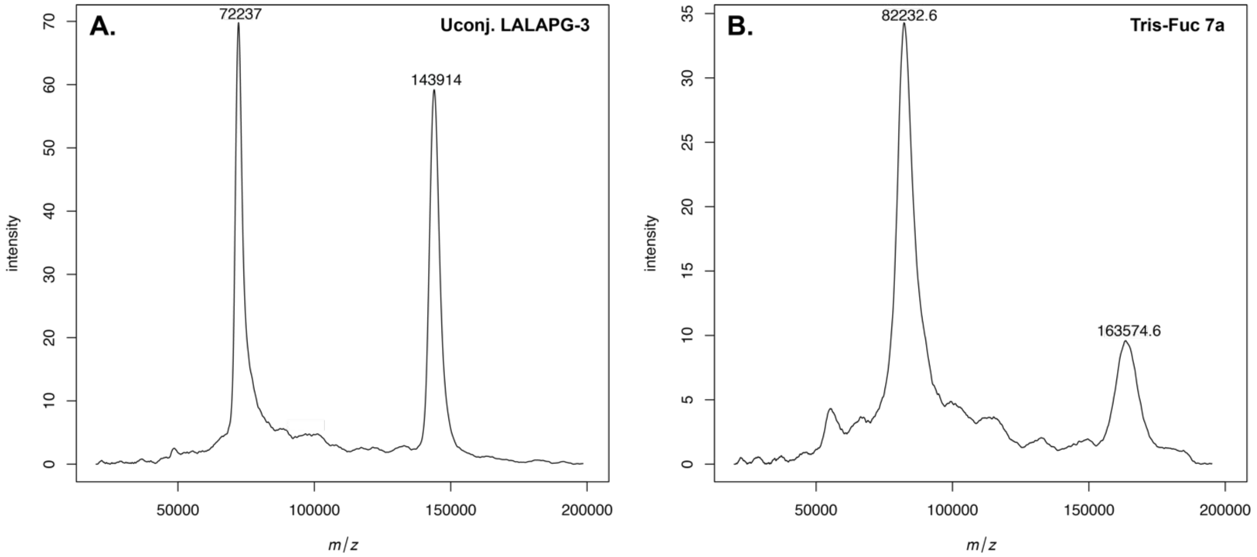


**Supplementary Figure S13.** MALDI-MS characterization of anti-SIRPα antibody conjugates. **A,** Combined spectrum of unconjugated LALAPG-3; **B,** Combined spectrum of tris-fucose conjugated anti-SIRPα antibody 3 (**7a**), where the average LAR is 16.

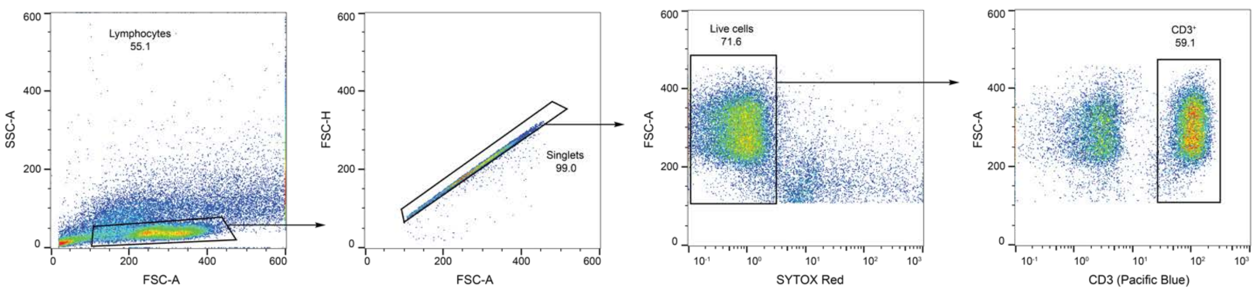


**Supplementary Figure S14.** Representative gating strategy for isolating T cells ($CD3^+$ cells) within healthy donor PBMCs. CD3 levels correspond to Pacific Blue, and Sytox Red was used for live/dead cell discrimination.

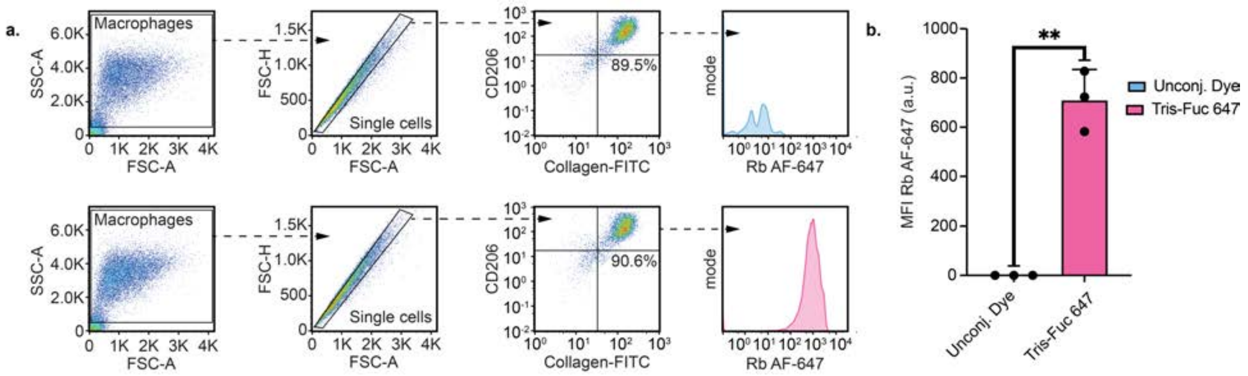


**Supplementary Figure S15. a,** Illustration of flow cytometry gating scheme for macrophage uptake for the various treatments and showing preservation of cell surface CD206 levels following co-treatment of 50 nM rabbit IgG-647 antibody with 25 nM of either unconjugated dye or tris-Fuc-647. **b,** Statistical analysis of M2-polarized macrophage uptake of anti-rabbit antibody uptake of tris-Fuc ligand. For **b**, error bars represent the SD from 3 independent experiments, *P* values were determined by Welch's two-tailed *t*-tests. Statistical significance was defined as $P<0.05$, and the asterisks * indicates $P<0.05$, ** indicates $P<0.01$, *** indicates $P<0.001$, and **** indicates $P<0.0001$.

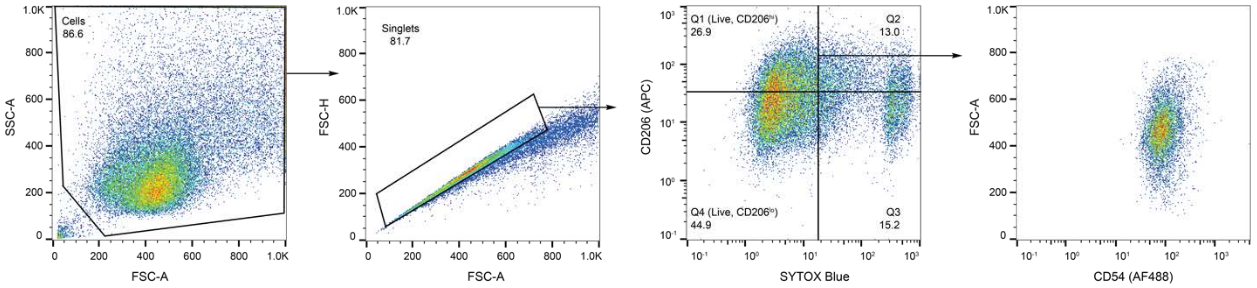


**Supplementary Figure S16**. Representative gating strategy for uptake and membrane protein internalization experiments. Data shown is for RAW264.7 cells treated with anti-CD54 antibody or TICTACs, gated for live, CD206$^{hi}$ cells. CD206 levels correspond to APC, and CD54 levels correspond to AF488. Sytox Blue was used for live/dead cell discrimination.

### Unprocessed Gels and Blot Images

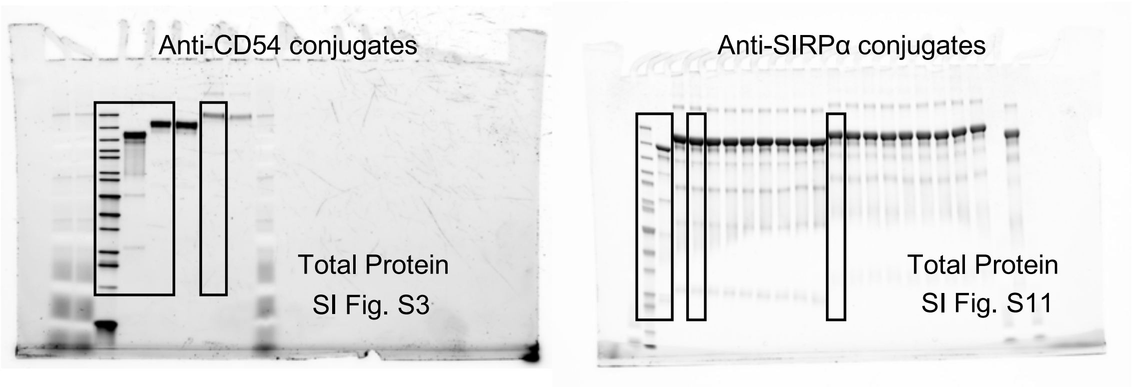


SDS-PAGE analysis of Anti-CD54 antibodies and Anti-SIRPa antibodies. Unconjugated (lane 1), NHS-PEG5-tris-alkyne conjugated (lanes 2-3 for left gel and lanes 2-10 for right gel), and tris-fucose conjugated (lanes 4-5 for left gel and lanes 11-19 for right gel). Total protein visualized by SYPRO Ruby stain. M, molecular weight marker.

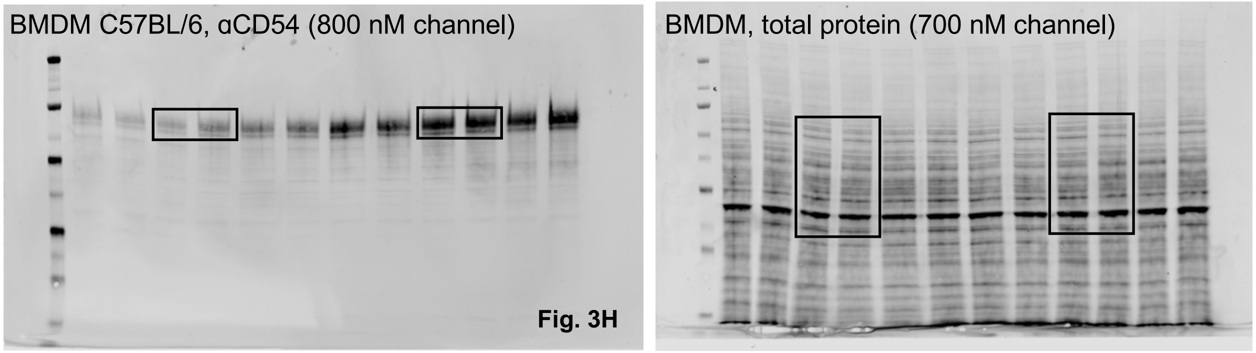


M2-polarized BMDMs from C57BL/6 mice were treated with 25 nM **4a**, **4b**, or unconjugated anti-CD54 antibody for 24 h. The total cell lysate was analyzed by Western blot. *Lanes from left to right*: 1–3: **4a** treatment (3 biological replicates); 4–6: **4b** treatment (3 biological replicates); 7–9: unconjugated anti-CD54 treatment (3 biological replicates); 10–12: No treatment (3 biological replicates).

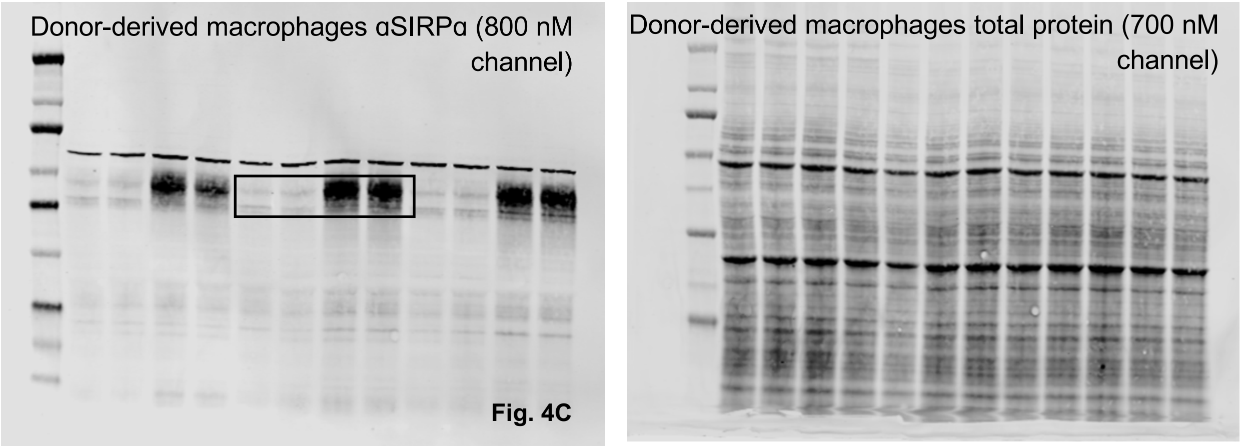


M2-polarized primary human macrophages from a representative donor were treated with 25 nM **5a**, **5b**, or unconjugated LALAPG-119 antibody for 24 h. *Lanes from left to right*: 1: **5a** treatment; 2: **5b** treatment; 3:

LALAPG-119 treatment; 4: No treatment; 5–12 are replicates (from the same donor) of 1–4 in the same order of treatment conditions.

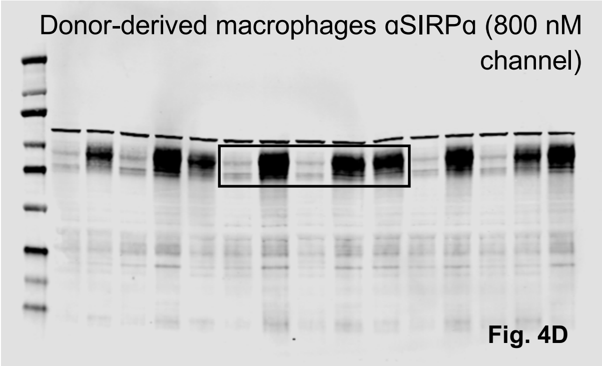


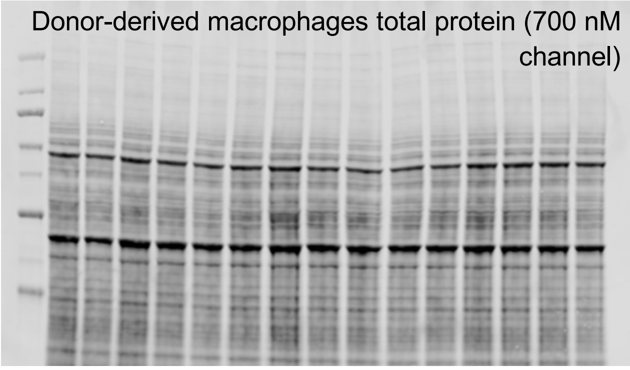


M2-polarized primary human macrophages from a representative donor were treated with 25 nM **6a**, **7a**, unconjugated LALAPG-136, or unconjugated LALAPG-3 antibody for 24 h. *Lanes from left to right*: 1: **6a** treatment; 2: LALAPG-136 treatment; 3: **7a** treatment; 4: LALAPG-3; 5: No treatment; 6–15 are replicates (from the same donor) of 1–5 in the same order of treatment conditions.

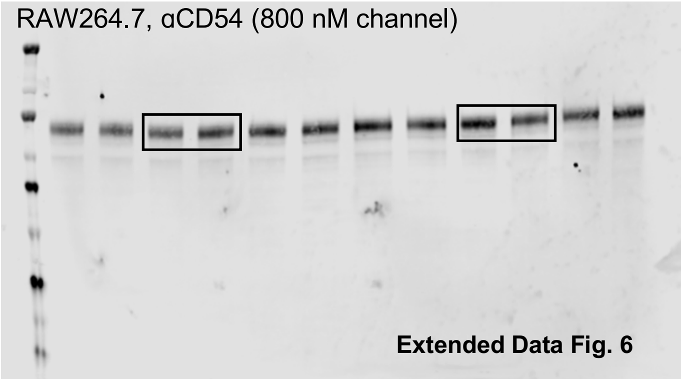


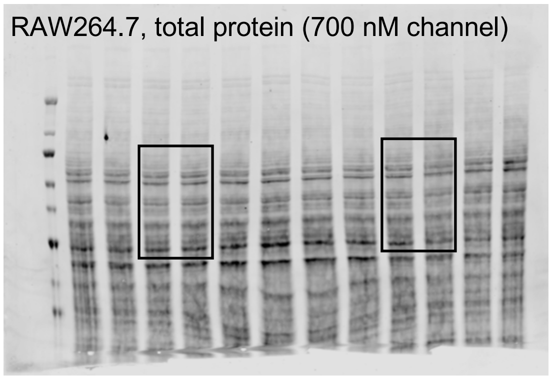


M2-polarized RAW264.7 cells were treated with 25 nM **4a**, **4b**, or unconjugated anti-CD54 antibody for 24 h. *Lanes from left to right*: 1–3: **4a** treatment (3 biological replicates); 4–6: **4b** treatment (3 biological replicates); 7–9: unconjugated anti-CD54 treatment (3 biological replicates); 10–12: No treatment (3 biological replicates).

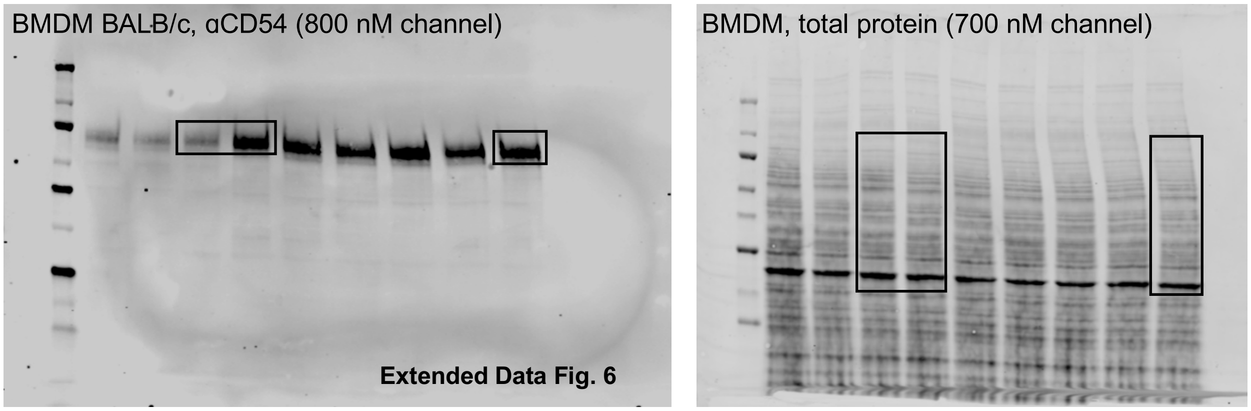


M2-polarized BMDMs from BALB/c mice were treated with 25 nM **4a** or unconjugated anti-CD54 antibody for 24 h. *Lanes from left to right*: 1–3: **4a** treatment (3 biological replicates); 4–6: unconjugated anti-CD54 treatment (3 biological replicates); 7–9: No treatment (3 biological replicates).

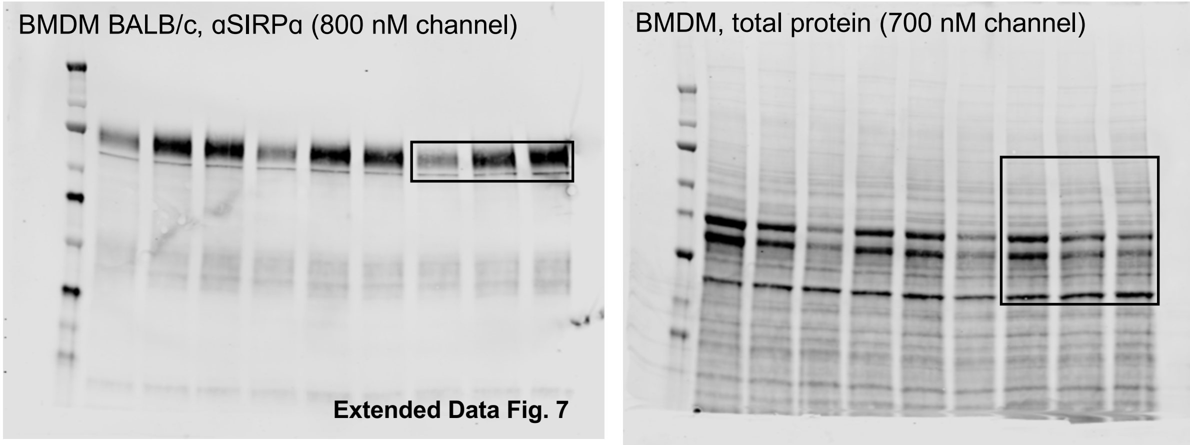


M2-polarized BMDMs from BALB/c mice were treated with 25 nM **6a** or unconjugated LALAPG-136 antibody for 24 h. *Lanes from left to right*: 1: **6a** treatment; 2: LALAPG-136 treatment; 3: No treatment; 4–9 are biological replicates of 1–3 in the same order of treatment conditions.

### Supplementary Note 1: General Methods and Instrumentation

Unless otherwise noted, all reactions were carried out in oven-dried glassware sealed with rubber septa under a nitrogen atmosphere with Teflon-coated magnetic stir bars. Reaction progress was monitored using thin layer chromatography on Millipore Sigma glass-backed TLC plates (250 µm thickness, F-254 indicator) and visualized with 254 nm UV light or stained by submersion in a basic potassium permanganate solution or 5% $H_2SO_4$ in methanol solution. Flash column chromatography was performed on a Biotage instrument using pre-packed silica gel columns. Anhydrous dimethylformamide (DMF), tetrahydrofuran (THF), and methylene chloride (DCM) were obtained from Sure/Seal bottles under an atmosphere of $N_2$. Deuterated solvents were purchased from Cambridge Isotope Laboratories. All other chemical reagents were purchased from Sigma Aldrich, Oakwood Chemicals, Conju-Probe, AxisPharm, Vector Laboratories, or Fischer Scientific and were used as directly received without further purification. Proton nuclear magnetic resonance ($^1H$ NMR) and carbon nuclear magnetic resonance ($^{13}C$ NMR) spectra were taken with Neo-500 Bruker spectrometer operating at 500 MHz at 25 °C. Chemical shifts are reported in parts per million (ppm) with reference to the appropriate residual solvent signal. $^1H$ NMR: $CDCl_3$ (δ: 7.26 ppm), DMSO-$d_6$ (δ: 2.50 ppm), MeOD (δ: 3.31 ppm), $D_2O$ (δ: 4.79 ppm). $^{13}C$NMR: MeOD (δ: 49.00). $^1H$ NMR multiplicities are reported as follows: s (singlet), d (doublet), t (triplet), q (quartet), sept (septet), m (multiplet).

*Liquid chromatography mass spectrometry (LC-MS) analysis.* Liquid chromatography mass spectrometry (LC-MS) analysis was performed on an Agilent 1260 Infinity II high performance liquid chromatography (HPLC) system with an Agilent 6125 single quadrupole LC/MSD. Samples were injected onto an Agilent InfinityLab Poroshell 120 Aq-C18 column (2.1 x 100 mm, 2.7 µm) using a gradient of 5% to 95% mobile phase B (0–1 min, 5%B; 1–10 min, 5–95%B; 10–12min, 95%B; 12–14 min, 95–5%B; mobile phase A set to 0.1% formic acid in water; mobile phase B set to 0.1% formic acid in acetonitrile). Flow rate was set to 700 µL/min. Data was collected and analyzed using OpenLab ChemStation software (Agilent Technologies).

*Preparative high performance liquid chromatography mass spectrometry (Prep-HPLC-MS) analysis*. Prep-HPLC-MS was performed on an Agilent 1290 Infinity II preparative high performance liquid chromatography (prep-HPLC) system coupled to an Agilent 6125 single quadrupole LC/MSD (Agilent Technologies). Samples were injected onto an Agilent 5 Prep 100Å C18 column (21.2 x 50 mm, 5 µm) using a gradient of 5% to 95% mobile phase B (0–0.5 min, 5%B; 0.5–6 min, 5%–60%B; 6–8 min, 60%–95%B; 8–11 min, 95%B; 11–12 min, 95%–5%B; mobile phase A set to 0.1% formic acid in water; mobile phase B set to 0.1% formic acid in acetonitrile). The preparative binary pump flow rate was set to 20 mL/min. The isocratic pump flow rate (for MSD) was set to 0.8 mL/min where the mobile phase was set to 0.1% formic acid in 20% water and 80% acetonitrile. Fractions were collected in 7 second increments throughout the run. Data was collected and analyzed using OpenLab ChemStation software (Agilent Technologies).

*High resolution mass spectrometry (HRMS) analysis.* HRMS data was acquired using an Agilent 1260 Infinity II high performance liquid chromatography (HPLC) system coupled to an Agilent

6230 time of flight (ToF) LC/MS (Agilent Technologies). Samples were injected onto an Agilent InfinityLab Poroshell 120 Aq-C18 column (2.1 x 100 mm, 2.7 μm) using a gradient of 5% to 100% mobile phase B (0–2 min, 5% B; 2–8 min, 5%–70% B; 8–10 min, 70%–100% B; 10–15 min, 100% B, 15–15.1 min, 100%–5% B; 17 min, 5% B; mobile phase A set to 0.2% formic acid in water; mobile phase B set to 0.2% formic acid in acetonitrile). Flow rate was set to 700 μL/min with a column temperature of 35 °C. Mass spectra were taken at a scan rate of 1 Hz over a 100–3200 *m/z* scan range with the ToF set to extended dynamic range. The mass spectrometer was operated using a dual Agilent jet stream (AJS) high-sensitivity ion source with the following instrument parameters: gas temperature (325 °C), drying gas (12 L/min), nebulizer (40 psi), sheath gas temperature (400 °C), sheath gas flow (12 L/min), VCap(3500 V), nozzle voltage (2000 V), fragmentor (20 V), skimmer (10 V), and Oct 1 RF Vpp (800 V). Mass spectra were analyzed and output from MassHunter software (Agilent Technologies).

### Supplementary Note 2: Synthesis and Characterization of Compounds

#### Propargyl 2,3,4,6-tetra-O-acetyl β-D-galactopyranoside (S1)

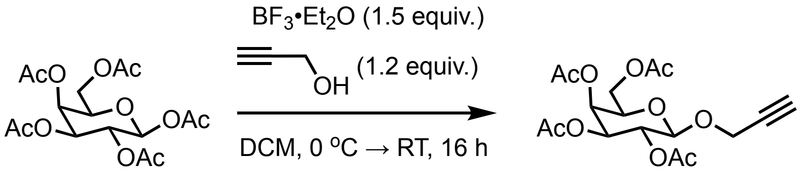


**S1**

β-D-galactose pentaacetate (2 g, 5.1 mmol, 1 equiv.) and propargyl alcohol (0.36 mL, 6.12 mmol, 1.2 equiv.) were combined in a flame-dried 50 mL Schlenk flask in 20 mL of anhydrous DCM. The mixture was cooled to 0 °C, and $BF_3 \cdot Et_2O$ (0.94 mL, 7.65 mmol, 1.5 equiv.) was added dropwise. The reaction was gradually warmed to RT and stirred overnight. The reaction was quenched with aq. $NaHCO_3$ (5 mL), stirred 20 min, diluted with water, and extracted with DCM (20 mL x 2). The combined organic phases were dried over magnesium sulfate, filtered, and concentrated *in vacuo*. The product was purified by column chromatography (30% EtOAc/hexanes) to yield **S1** as a clear oil. The $^1$H NMR data matched that previously reported in the literature[2]. $^1$H NMR (500 MHz, Chloroform-*d*) δ 5.40 (dd, *J* = 3.5, 1.1 Hz, 1H), 5.22 (dd, *J* = 10.4, 7.9 Hz, 1H), 5.06 (dd, *J* = 10.4, 3.4 Hz, 1H), 4.74 (d, *J* = 8.0 Hz, 1H), 4.38 (d, *J* = 2.3 Hz, 2H), 4.22 – 4.09 (m, 2H), 3.94 (td, *J* = 6.7, 1.2 Hz, 1H), 2.46 (t, *J* = 2.4 Hz, 1H), 2.15 (s, 3H), 2.07 (s, 3H), 2.04 (s, 3H), 1.99 (s, 3H).

#### Propargyl β-D-galactopyranoside (S2)

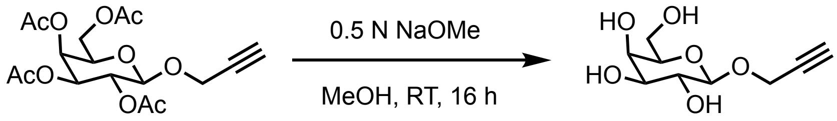


**S2**

To a solution of compound **S1** in anhydrous MeOH (10 mL) was added 0.5 N NaOMe solution (20 mL). The reaction was stirred at RT overnight and quenched with Amberlite IR-120 resin ($H^+$ form), filtered, then concentrated to obtain the crude product, which was a semi-solid/viscous oil. The crude product was triturated in pentane/$Et_2O$, filtered, and dried under high vacuum for 1 h to yield **S2** as a crystalline white solid ($R_f$ = 0.25 in 3:1 $CHCl_3$: MeOH). The $^1$H NMR data matched that previously reported in the literature[2]. $^1$H NMR (500 MHz, MeOD/Tol 1:1) δ 4.48 (d, *J* = 7.7 Hz, 1H), 4.44 (dd, *J* = 4.0, 2.5 Hz, 2H), 3.93 – 3.87 (m, 2H), 3.84 (dd, *J* = 11.4, 5.4 Hz, 1H), 3.72 (dd, *J* = 9.6, 7.8 Hz, 1H), 3.56 (dd, *J* = 9.7, 3.4 Hz, 1H), 3.50 (t, *J* = 6.1 Hz, 1H), 2.67 (t, *J* = 2.5 Hz, 1H).

**Sodium propargyl 3-sulfonato-β-D-galactopyranoside (S3)**

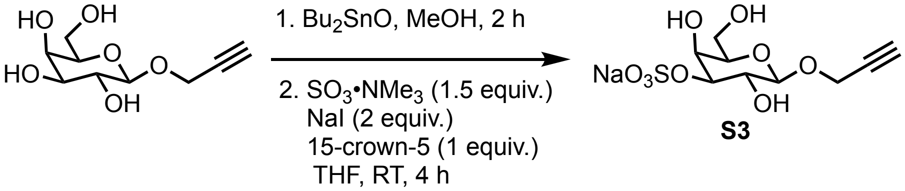


Compound **S2** (324 mg, 1.48 mmol, 1 equiv.) and dibutyltin oxide (555 mg, 2.23 mmol, 1.5 equiv.) were dissolved in anhydrous MeOH (30 mL) and the reaction was heated to reflux for 2 h. The solvent was removed under reduced pressure, and the residue was taken up in anhydrous THF (20 mL). Sulfur trioxide trimethylamine complex (355 mg, 2.23 mmol, 1.5 equiv.) was first added, then NaI (445 mg, 2.97 mmol, 2 equiv.), then 15-crown-5 (0.3 mL, 1.48 mmol, 1 equiv.), and the reaction was stirred at RT for 4 h. The reaction was monitored with TLC (3:1 $CHCl_3$: MeOH). When the reaction was complete, $NaHCO_3$ (5 mL) and 5% Na2S2O3 (5 mL) were added. The mixture was concentrated *in vacuo*, then directly adsorbed onto silica in MeOH. The crude mixture was purified by column chromatography ($CHCl_3$, then $CHCl_3$: MeOH 9:1, $CHCl_3$: MeOH 5:1, then 1:1). Earlier impurities elute with 9:1 and 5:1. **S3** was isolated as a white foam (404 mg, 85% yield). The $^1H$ NMR data matched that previously reported in the literature[2]. $^1H$ NMR (500 MHz, MeOD/Tol 1:1) δ 4.50 (d, $J$ = 7.7 Hz, 1H), 4.38 – 4.30 (m, 4H), 3.87 – 3.72 (m, 3H), 3.49 (t, $J$ = 6.0 Hz, 1H), 2.61 (q, $J$ = 1.8 Hz, 1H).

**Synthesis of Propargyl α-D-mannose (S4)**

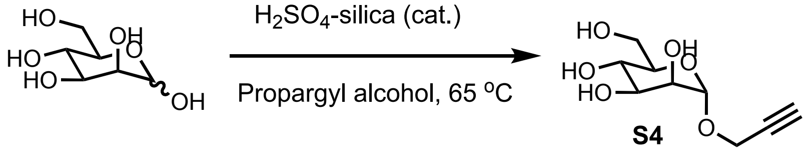


To mannose (1 g, 1 equiv.) in neat propargyl alcohol (1.6 mL) was added $H_2SO_4$-silica* (27.8 mg, cat.). The reaction was heated to 65 °C, during the course of which the mixture became homogeneous. The crude mixture was first eluted through a short silica column with DCM to remove excess propargyl alcohol, then purified by column chromatography (10% MeOH in DCM) to isolate the α-anomer **S4** as a white solid. The $^1H$ NMR data matched that previously reported in the literature.[3] $^1H$ NMR (500 MHz, MeOD) δ 4.96 (d, $J$ = 1.7 Hz, 1H), 4.27 (d, $J$ = 2.4 Hz, 2H), 3.86 – 3.78 (m, 2H), 3.73 – 3.59 (m, 3H), 3.51 (ddd, $J$ = 10.8, 5.1, 2.1 Hz, 1H), 2.85 (t, $J$ = 2.4 Hz, 1H).

**$H_2SO_4$-slica was prepared as the following:*

10 g silica was added to diethyl ether (50 mL) and mixed to form a slurry. To this mixture was added concentrated $H_2SO_4$ (3 mL), and the mixture was shaken/stirred for 5 min. The solvent was removed under reduced pressure until silica was free flowing, then dried in the oven for at least 3 h.

**Amino-PEG$_4$-tris-α-L-fucose (S5)**

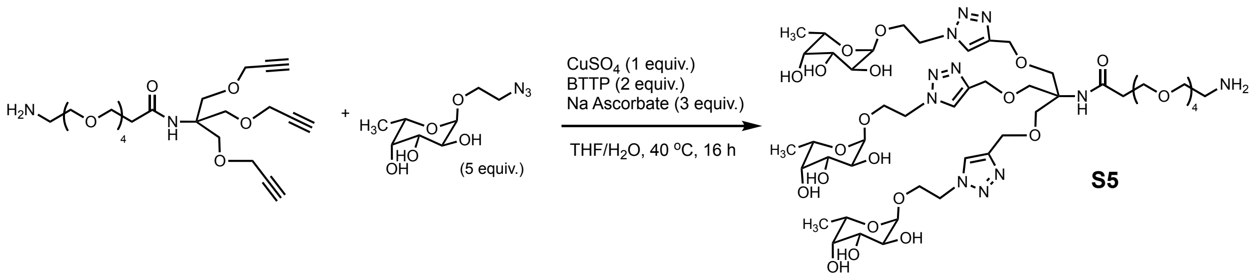


To a 1-dram vial equipped with a magnetic stir bar was added amino-PEG$_4$-tris-alkyne (10 mg, 0.02 mmol, 1 equiv.) and 2-azidoethyl α-L-fucopyranoside (23.3 mg, 0.14 mmol, 5 equiv.) as a solution in 0.5 mL of degassed 1:1 THF/ddH$_2$O. In a separate one-dram vial, $CuSO_4$ (5 mg, 0.02 mmol, 1 equiv.), BTTP (17.22 mg, 0.04 mmol, 2 equiv.), and sodium ascorbate (11.9 mg, 0.06 mmol, 3 equiv.) were combined in 0.2 mL of ddH$_2$O, then added to the reaction mixture. The solution was heated to 40 °C and stirred for 16 h. The mixture was purified by preparative HPLC using a 0-30% acetonitrile gradient over 11 min, and the product was isolated as a white solid after lyophilization (15.5 mg, 65% yield). $^{1}$H NMR (500 MHz, MeOD) δ 8.11 (s, 3H), 4.76 (d, *J* = 3.8 Hz, 3H), 4.64 (dddd, *J* = 18.3, 14.8, 6.0, 3.5 Hz, 7H), 4.58 (s, 6H), 4.06 (ddd, *J* = 11.1, 7.8, 3.5 Hz, 3H), 3.83 (ddd, *J* = 10.8, 5.2, 3.5 Hz, 4H), 3.77 (s, 6H), 3.75 – 3.73 (m, 3H), 3.72 (d, *J* = 3.8 Hz, 3H), 3.71 – 3.67 (m, 5H), 3.66 (d, *J* = 3.3 Hz, 2H), 3.65 – 3.61 (m, 8H), 3.59 (s, 5H), 3.57 (dd, *J* = 3.3, 1.3 Hz, 4H), 3.43 – 3.38 (m, 3H), 3.15 (d, *J* = 5.1 Hz, 2H), 2.48 (t, *J* = 6.0 Hz, 2H), 1.10 (d, *J* = 6.6 Hz, 9H). $^{13}$C NMR (126 MHz, MeOD) δ 174.24, 145.78, 126.35, 100.44, 73.44, 71.54, 71.25, 71.22, 69.82, 69.50, 67.71, 67.13, 65.33, 51.29, 40.58, 37.91, 16.66. HRMS (ESI-TOF) Calcd for $C_{48}H_{84}N_{11}O_{23}$ [M+H]$^+$: 1182.5736, Found: 1181.5700.

**Tris-fucose-647** (**2**)

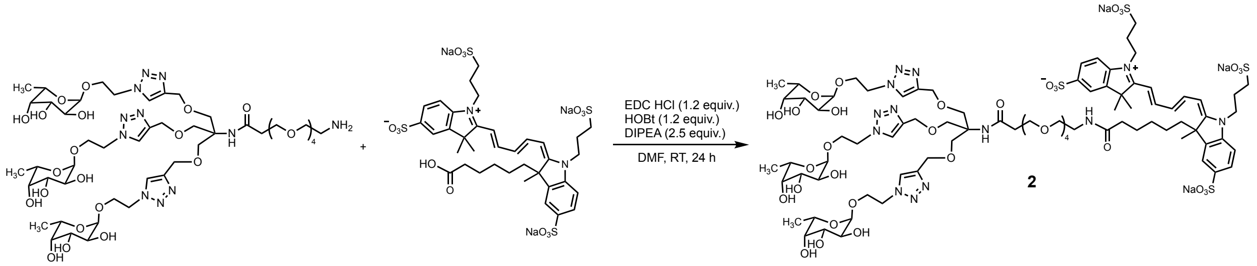


To a 1-dram vial equipped with a magnetic stir bar was added **S5** (3.62 mg, 0.0031 mmol, 1 equiv.), APDye Fluor 647 acid (3 mg, 0.0032 mmol, 1 equiv.), and 1-hydroxy-7-azabenzotriazole (0.52 mg, 0.0038 mmol, 1.2 equiv.) as a solution in 0.2 mL DMF. To this mixture was added 1-ethyl-3-(3-dimethylaminopropyl)carbodiimide hydrochloride (0.73 mg, 0.0038 mmol, 1.2 equiv.) and diisopropylethylamine (1.33 µL, 0.0077 mmol, 2.5 equiv.). The mixture was stirred for 24 h at room temperature. The crude mixture was purified by prep-HPLC using a 5-95% acetonitrile gradient over 11 min, and the product was isolated as a bright blue solid after lyophilization. HRMS (ESI-TOF) Calcd for $C_{84}H_{124}N_{13}Na_2O_{36}S_4$ [M-Na]$^-$: 2064.6911, Found: 2064.6400.

### $^1$H and $^{13}$C NMR spectra

### Compound S5: 1H NMR

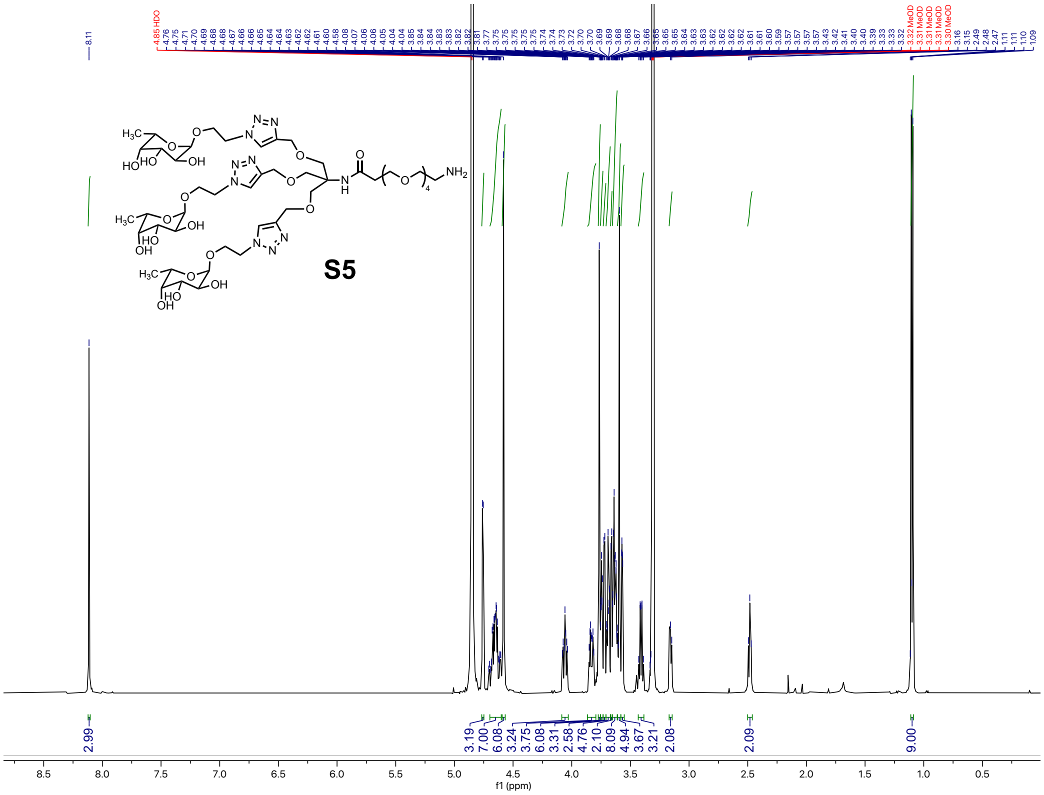


### Compound S5: 13C NMR

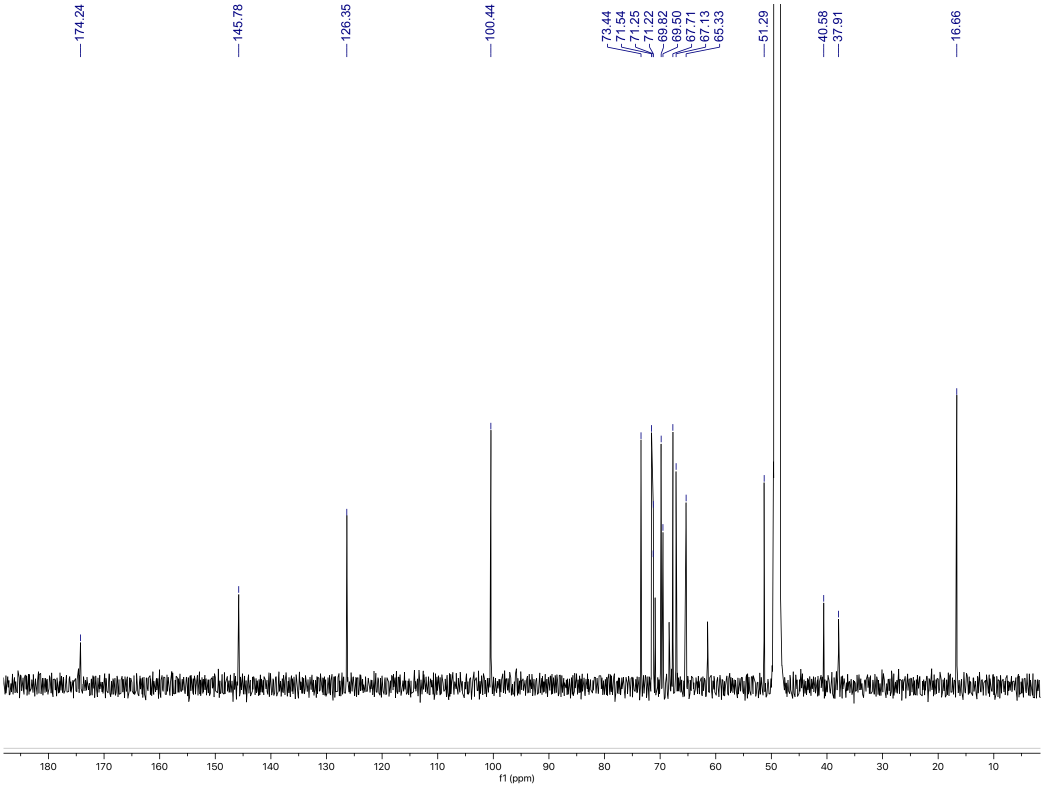

### Mass spectrometry characterization

#### Tris-fucose-647 (2): ESI-MS

**A.** TIC (ESI$^-$)

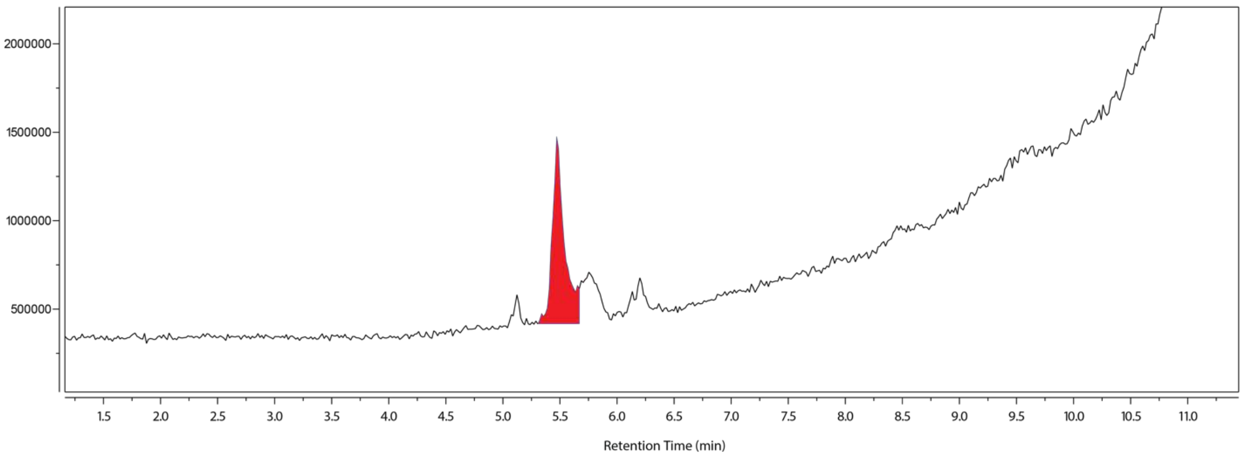


**B.** HRMS (5.42-5.60 min)

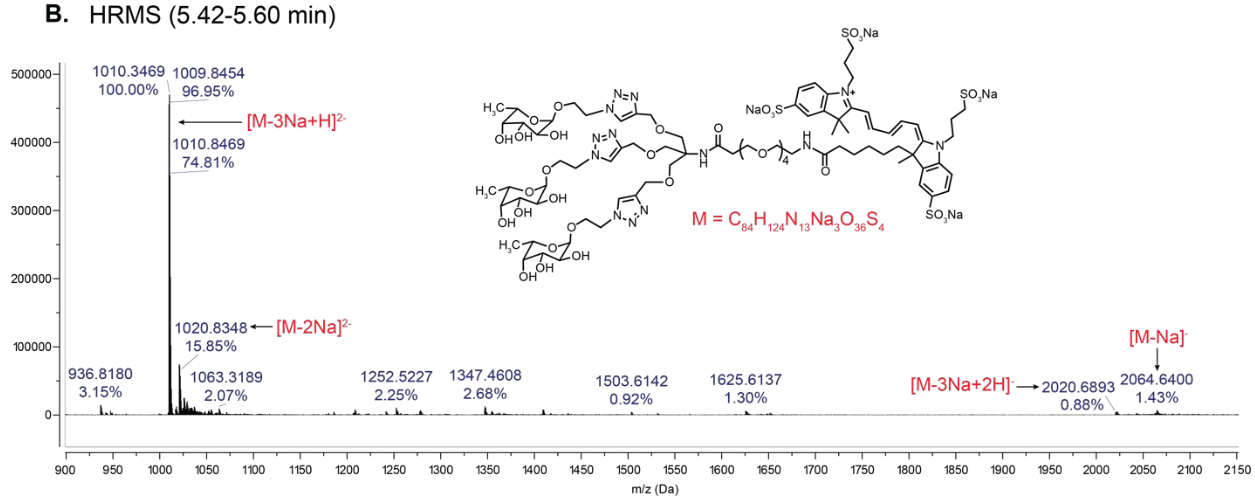


**A,** Total ion chromatogram (TIC) of compound **2** measured in the negative ion mode, where the peak of interest is highlighted in red. **B,** High resolution mass spectrometry (HRMS) data for the peak eluting at 5.42-5.60 min, where relevant peaks are annotated in red. The parent monoisotopic mass M was set to $C_{84}H_{124}N_{13}Na_3O_{36}S_4$.

### Supplementary Note 3

#### Amino Acid Sequences for LALAPG-119, LALAPG-136, and LALAPG-3

| Name | Amino Acid Sequence |
|---|---|
| LALAPG-119 heavy chain with signal peptide (signal peptide in green) | MHSSALLCCLVLLTGVRADVQLLESGGGVVQPGGSLRLSCAASGFS<br>FSNFAMTWVRQAPGEGLEWVSTIGSGDTYYADSVKGRFTISRDNSK<br>NTLYLQMNSLRAEDTAVYYCAKDSTVSWSGDFFDYWGQGTLVTVS<br>SASTKGPSVFPLAPSSKSTSGGTAALGCLVKDYFPEPVTVSWNSGA<br>LTSGVHTFPAVLQSSGLYSLSSVVTVPSSSLGTQTYICNVNHKPSNT<br>KVDKKVEPKSCDKTHTCPPCPAPEAAGGPSVFLFPPKPKDTLMISRT<br>PEVTCVVVDVSHEDPEVKFNWYVDGVEVHNAKTKPREEQYNSTYR<br>VVSVLTVLHQDWLNGKEYKCKVSNKALGAPIEKTISKAKGQPREPQV<br>YTLPPSRDELTKNQVSLTCLVKGFYPSDIAVEWESNGQPENNYKTT<br>PPVLDSDGSFFLYSKLTVDKSRWQQGNVFSCSVMHEALHNHYTQK<br>SLSLSPGK |
| LALAPG-119 light chain with signal peptide (signal peptide in green) | MHSSALLCCLVLLTGVRAEIVLTQSPATLSVSPGERATFSCRASQNV<br>KNDLAWYQQRPGQAPRLLIYAARIRETGIPERFSGSGSGTEFTLTITS<br>LQSEDFAVYYCQQYYDWPPFTFGGGTKVEIKRTVAAPSVFIFPPSDE<br>QLKSGTASVVCLLNNFYPREAKVQWKVDNALQSGNSQESVTEQDS<br>KDSTYSLSSTLTLSKADYEKHKVYACEVTHQGLSSPVTKSFNRGEC |
| LALAPG-136 heavy chain with signal peptide (signal peptide in green) | MHSSALLCCLVLLTGVRADVQLVESGGGVVRPGESLRLSCAASGFT<br>FSSYDMNWVRQAPGEGLEWVSLISGSGEIIYYADSVKGRFTISRDNS<br>KNTLYLQMNSLRAEDTAVYYCAKENNRYRFFDDWGQGTLVTVSSA<br>STKGPSVFPLAPSSKSTSGGTAALGCLVKDYFPEPVTVSWNSGALT<br>SGVHTFPAVLQSSGLYSLSSVVTVPSSSLGTQTYICNVNHKPSNTKV<br>DKKVEPKSCDKTHTCPPCPAPEAAGGPSVFLFPPKPKDTLMISRTPE<br>VTCVVVDVSHEDPEVKFNWYVDGVEVHNAKTKPREEQYNSTYRVV<br>SVLTVLHQDWLNGKEYKCKVSNKALGAPIEKTISKAKGQPREPQVYT<br>LPPSRDELTKNQVSLTCLVKGFYPSDIAVEWESNGQPENNYKTTPP<br>VLDSDGSFFLYSKLTVDKSRWQQGNVFSCSVMHEALHNHYTQKSL<br>SLSPGK |
| LALAPG-136 light chain with signal peptide (signal peptide in green) | MHSSALLCCLVLLTGVRAETVLTQSPGTLTLSPGERATLTCRASQSV<br>YTYLAWYQEKPGQAPRLLIYGASSRATGIPDRFSGSGSGTEFTLTIS<br>SLQSEDFAVYYCQQYYDRPPLTFGGGTKVEIKRTVAAPSVFIFPPSD<br>EQLKSGTASVVCLLNNFYPREAKVQWKVDNALQSGNSQESVTEQD<br>SKDSTYSLSSTLTLSKADYEKHKVYACEVTHQGLSSPVTKSFNRGE<br>C |
| LALAPG-3 heavy chain with signal peptide (signal peptide in green) | MHSSALLCCLVLLTGVRAAVTLDESGGGLQTPGGALSLVCKASGFIF<br>SDYGMNWVRQAPGKGLEFVAQITSGSRTYYGAAVKGRATISRDNR<br>QSTVKLQLNNLRAEDTGIYFCARDFGSGVGSIDAWGNGTEVIVSSAS<br>TKGPSVFPLAPSSKSTSGGTAALGCLVKDYFPEPVTVSWNSGALTS<br>GVHTFPAVLQSSGLYSLSSVVTVPSSSLGTQTYICNVNHKPSNTKVD<br>KKVEPKSCDKTHTCPPCPAPEAAGGPSVFLFPPKPKDTLMISRTPEV<br>TCVVVDVSHEDPEVKFNWYVDGVEVHNAKTKPREEQYNSTYRVVS<br>VLTVLHQDWLNGKEYKCKVSNKALGAPIEKTISKAKGQPREPQVYTL<br>PPSRDELTKNQVSLTCLVKGFYPSDIAVEWESNGQPENNYKTTPPV<br>LDSDGSFFLYSKLTVDKSRWQQGNVFSCSVMHEALHNHYTQKSLSL<br>SPGK |
| LALAPG-3 light chain with signal peptide (signal peptide in green) | MHSSALLCCLVLLTGVRAALTQPASVSANLGGTVKITCSGSRGRYG<br>WYQQRSPGSAPVTVIYRDNQRPSNIPSRFSSSTSGSTSTLTITGVQA<br>DDESVYFCGSYDGSIDIFGAGTTLTVLRTVAAPSVFIFPPSDEQLKSG<br>TASVVCLLNNFYPREAKVQWKVDNALQSGNSQESVTEQDSKDSTY<br>SLSSTLTLSKADYEKHKVYACEVTHQGLSSPVTKSFNRGEC |

### DNA Sequences for LALAPG-119, LALAPG-136, and LALAPG-3

| Name | DNA Sequence |
|---|---|
| LALAPG-119 heavy chain with signal peptide | GCGGCCGCAAACTACAAGACAGACTTGCAAAAGAAGGCATGCAC AGCTCAGCACTGCTCTGTTGCCTGGTCCTCCTGACTGGGGTGAG GGCCGATGTGCAGCTGTTGGAGTCCGGGGGAGGAGTTGTGCAG CCTGGAGGAAGTTTGAGGTTGTCCTGTGCCGCCTCCGGGTTTTC CTTTTCCAACTTTGCCATGACCTGGGTGAGGCAGGCTCCCGGAG AAGGATTGGAGTGGGTGAGCACTATTGGCTCCGGAGATACCTAC TACGCCGACTCCGTTAAGGGCAGGTTCACTATCTCTCGGGACAA CAGCAAAAACACCCTGTACCTGCAGATGAACTCCCTGAGGGCCG AGGACACCGCCGTGTACTACTGCGCCAAGGACTCCACCGTGTCC TGGTCCGGCGACTTCTTCGACTACTGGGGCCAGGGCACCCTGGT GACCGTGTCCTCCGCTAGCACCAAGGGCCCATCGGTCTTCCCCC TGGCACCCTCCTCCAAGAGCACCTCTGGGGGCACAGCGGCCCT GGGCTGCCTGGTCAAGGACTACTTCCCCGAACCGGTGACGGTGT CGTGGAACTCAGGCGCCCTGACCAGCGGCGTGCACACCTTCCC GGCTGTCCTACAGTCCTCAGGACTCTACTCCCTCAGCAGCGTGG TGACCGTGCCCTCCAGCAGCTTGGGCACCCAGACCTACATCTGC AACGTGAATCACAAGCCCAGCAACACCAAGGTGGACAAGAAAGT TGAGCCCAAATCTTGTGACAAAACTCACACATGCCCACCGTGCCC AGCACCTGAAGCCGCAGGGGGACCGTCAGTCTTCCTCTTCCCCC CAAAACCCAAGGACACCCTCATGATCTCCCGGACCCCCGAGGTC ACATGCGTGGTGGTGGACGTGAGCCACGAAGACCCTGAGGTCAA GTTCAACTGGTACGTGGACGGCGTGGAGGTGCATAATGCCAAGA CAAAGCCGCGGGAGGAGCAGTACAACAGCACGTACCGTGTGGT CAGCGTCCTCACCGTCCTGCACCAGGACTGGCTGAATGGCAAGG AGTACAAGTGCAAGGTCTCCAACAAAGCCCTCGGAGCCCCCATC GAGAAAACCATCTCCAAAGCCAAAGGGCAGCCCCGAGAACCACA GGTGTACACCCTGCCCCCATCCCGGGACGAGCTGACCAAGAACC AGGTCAGCCTGACCTGCCTGGTCAAAGGCTTCTATCCCAGCGAC ATCGCCGTGGAGTGGGAGAGCAATGGGCAGCCGGAGAACAACT ACAAGACCACGCCTCCCGTGCTGGACTCCGACGGCTCCTTCTTC CTCTACAGCAAGCTCACCGTGGACAAGAGCAGGTGGCAGCAGG GGAACGTCTTCTCATGCTCCGTGATGCATGAGGCTCTGCACAAC CACTACACGCAGAAGAGCCTCTCCCTGTCTCCGGGTAAATGATTC TAGA |
| LALAPG-119 light chain with signal peptide | GCGGCCGCAAACTACAAGACAGACTTGCAAAAGAAGGCATGCAC AGCTCAGCACTGCTCTGTTGCCTGGTCCTCCTGACTGGGGTGAG GGCCGAAATCGTGCTGACCCAGTCCCCCGCTACCTTGAGTGTGT CCCCTGGAGAGAGAGCCACCTTCAGCTGTAGGGCTTCCCAGAAC GTGAAAAACGACCTGGCCTGGTACCAGCAGAGGCCCGGCCAGG CCCCCAGGCTGCTGATCTACGCCGCCAGGATCAGGGAGACCGG CATCCCCGAGAGGTTCTCCGGCTCCGGCTCCGGCACCGAGTTCA CCCTGACCATCACCTCCCTGCAGTCCGAGGACTTCGCCGTGTAC TACTGCCAGCAGTACTACGACTGGCCCCCCTTCACCTTCGGCGG CGGCACCAAGGTGGAGATCAAGCGTACGGTGGCTGCACCATCTG TCTTCATCTTCCCGCCATCTGATGAGCAGTTGAAATCTGGAACTG CCTCTGTTGTGTGCCTGCTGAATAACTTCTATCCCAGAGAGGCCA AAGTACAGTGGAAGGTGGATAACGCCCTCCAATCGGGTAACTCC CAGGAGAGTGTCACAGAGCAGGACAGCAAGGACAGCACCTACA GCCTCAGCAGCACCCTGACGCTGAGCAAAGCAGACTACGAGAAA CACAAAGTCTACGCCTGCGAAGTCACCCATCAGGGCCTGAGTTC GCCCGTCACAAAGAGCTTCAACAGGGGAGAGTGTTGATTCTAGA |

| | |
|---|---|
| LALAPG-136 heavy chain with signal peptide | GCGGCCGCAAACTACAAGACAGACTTGCAAAAGAAGGCATGCAC<br>AGCTCAGCACTGCTCTGTTGCCTGGTCCTCCTGACTGGGGTGAG<br>GGCCGATGTGCAGCTGGTGGAGTCCGGAGGAGGAGTTGTGAGG<br>CCTGGAGAGTCCCTGAGGTTGTCCTGTGCCGCTTCCGGATTTAC<br>CTTTTCCTCCTACGACATGAACTGGGTGAGGCAGGCTCCCGGAG<br>AGGGATTGGAATGGGTGAGCTTGATCTCCGGCAGTGGGGAGATC<br>ATCTATTACGCTGACTCTGTTAAGGGCCGGTTCACCATCTCCAGG<br>GACAACAGCAAGAACACCCTGTACTTGCAGATGAACTCCTTGCG<br>GGCCGAGGACACCGCTGTTTATTATTGCGCCAAGGAGAACAACA<br>GGTATAGGTTTTTCGACGACTGGGGCCAGGGAACCCTGGTTACA<br>GTGTCCTCCGCTAGCACCAAGGGCCCATCGGTCTTCCCCCTGGC<br>ACCCTCCTCCAAGAGCACCTCTGGGGGCACAGCGGCCCTGGGC<br>TGCCTGGTCAAGGACTACTTCCCCGAACCGGTGACGGTGTCGTG<br>GAACTCAGGCGCCCTGACCAGCGGCGTGCACACCTTCCCGGCT<br>GTCCTACAGTCCTCAGGACTCTACTCCCTCAGCAGCGTGGTGAC<br>CGTGCCCTCCAGCAGCTTGGGCACCCAGACCTACATCTGCAACG<br>TGAATCACAAGCCCAGCAACACCAAGGTGGACAAGAAAGTTGAG<br>CCCAAATCTTGTGACAAAACTCACACATGCCCACCGTGCCCAGCA<br>CCTGAAGCCGCAGGGGGACCGTCAGTCTTCCTCTTCCCCCCAAA<br>ACCCAAGGACACCCTCATGATCTCCCGGACCCCCGAGGTCACAT<br>GCGTGGTGGTGGACGTGAGCCACGAAGACCCTGAGGTCAAGTT<br>CAACTGGTACGTGGACGGCGTGGAGGTGCATAATGCCAAGACAA<br>AGCCGCGGGAGGAGCAGTACAACAGCACGTACCGTGTGGTCAG<br>CGTCCTCACCGTCCTGCACCAGGACTGGCTGAATGGCAAGGAGT<br>ACAAGTGCAAGGTCTCCAACAAAGCCCTCGGAGCCCCCATCGAG<br>AAAACCATCTCCAAAGCCAAAGGGCAGCCCCGAGAACCACAGGT<br>GTACACCCTGCCCCCATCCCGGGACGAGCTGACCAAGAACCAG<br>GTCAGCCTGACCTGCCTGGTCAAAGGCTTCTATCCCAGCGACAT<br>CGCCGTGGAGTGGGAGAGCAATGGGCAGCCGGAGAACAACTAC<br>AAGACCACGCCTCCCGTGCTGGACTCCGACGGCTCCTTCTTCCT<br>CTACAGCAAGCTCACCGTGGACAAGAGCAGGTGGCAGCAGGGG<br>AACGTCTTCTCATGCTCCGTGATGCATGAGGCTCTGCACAACCAC<br>TACACGCAGAAGAGCCTCTCCCTGTCTCCGGGTAAATGATTCTAG<br>A |
| LALAPG-136 light chain with signal peptide | GCGGCCGCAAACTACAAGACAGACTTGCAAAAGAAGGCATGCAC<br>AGCTCAGCACTGCTCTGTTGCCTGGTCCTCCTGACTGGGGTGAG<br>GGCCGAAACAGTGCTGACCCAGTCCCCCGGAACCTTGACATTGA<br>GCCCCGGAGAAAGGGCTACCCTGACCTGTAGGGCCTCCCAGTCT<br>GTGTACACCTACCTGGCCTGGTACCAGGAGAAGCCCGGACAGG<br>CTCCTCGATTGCTCATCTACGGCGCCTCCAGCAGGGCTACAGGA<br>ATCCCTGATAGGTTCTCCGGTTCCGGATCTGGCACCGAGTTCAC<br>CCTGACCATCTCCTCCCTGCAGAGCGAGGATTTCGCCGTCTACT<br>ACTGCCAGCAGTACTACGACCGTCCCCCCTTGACCTTTGGCGGA<br>GGAACAAAGGTTGAGATCAAACGTACGGTGGCTGCACCATCTGT<br>CTTCATCTTCCCGCCATCTGATGAGCAGTTGAAATCTGGAACTGC<br>CTCTGTTGTGTGCCTGCTGAATAACTTCTATCCCAGAGAGGCCAA<br>AGTACAGTGGAAGGTGGATAACGCCCTCCAATCGGGTAACTCCC<br>AGGAGAGTGTCACAGAGCAGGACAGCAAGGACAGCACCTACAG<br>CCTCAGCAGCACCCTGACGCTGAGCAAAGCAGACTACGAGAAAC<br>ACAAAGTCTACGCCTGCGAAGTCACCCATCAGGGCCTGAGTTCG<br>CCCGTCACAAAGAGCTTCAACAGGGGAGAGTGTTGATTCTAGA |
| LALAPG-3 heavy chain with signal peptide | GCGGCCGCAAACTACAAGACAGACTTGCAAAAGAAGGCATGCAC<br>AGCTCAGCACTGCTCTGTTGCCTGGTCCTCCTGACTGGGGTGAG<br>GGCCGCTGTTACCCTGGATGAGTCCGGAGGAGGATTGCAGACC<br>CCCGGAGGAGCTTTGTCCCTGGTTTGTAAGGCCTCTGGATTTATT<br>TTTTCTGATTATGGAATGAACTGGGTGAGGCAGGCCCCCGGAAA |

| | |
|---|---|
| | GGGATTGGAATTTGTGGCTCAGATCACCTCCGGATCTCGTACCTA<br>CTACGGCGCCGCAGTTAAGGGACGGGCTACTATCAGCAGGGAC<br>AACAGGCAGAGTACAGTGAAACTCCAGCTGAATAACCTGAGGGC<br>CGAAGACACCGGCATCTATTTCTGCGCCAGGGACTTTGGATCTG<br>GGGTGGGAAGCATTGACGCCTGGGGAAACGGAACAGAAGTGAT<br>CGTGAGCTCCGCTAGCACCAAGGGCCCATCGGTCTTCCCCCTGG<br>CACCCTCCTCCAAGAGCACCTCTGGGGGCACAGCGGCCCTGGG<br>CTGCCTGGTCAAGGACTACTTCCCCGAACCGGTGACGGTGTCGT<br>GGAACTCAGGCGCCCTGACCAGCGGCGTGCACACCTTCCCGGC<br>TGTCCTACAGTCCTCAGGACTCTACTCCCTCAGCAGCGTGGTGA<br>CCGTGCCCTCCAGCAGCTTGGGCACCCAGACCTACATCTGCAAC<br>GTGAATCACAAGCCCAGCAACACCAAGGTGGACAAGAAAGTTGA<br>GCCCAAATCTTGTGACAAAACTCACACATGCCCACCGTGCCCAG<br>CACCTGAAGCCGCAGGGGGACCGTCAGTCTTCCTCTTCCCCCCA<br>AAACCCAAGGACACCCTCATGATCTCCCGGACCCCCGAGGTCAC<br>ATGCGTGGTGGTGGACGTGAGCCACGAAGACCCTGAGGTCAAGT<br>TCAACTGGTACGTGGACGGCGTGGAGGTGCATAATGCCAAGACA<br>AAGCCGCGGGAGGAGCAGTACAACAGCACGTACCGTGTGGTCA<br>GCGTCCTCACCGTCCTGCACCAGGACTGGCTGAATGGCAAGGAG<br>TACAAGTGCAAGGTCTCCAACAAAGCCCTCGGAGCCCCCATCGA<br>GAAAACCATCTCCAAAGCCAAAGGGCAGCCCCGAGAACCACAGG<br>TGTACACCCTGCCCCCATCCCGGGACGAGCTGACCAAGAACCAG<br>GTCAGCCTGACCTGCCTGGTCAAAGGCTTCTATCCCAGCGACAT<br>CGCCGTGGAGTGGGAGAGCAATGGGCAGCCGGAGAACAACTAC<br>AAGACCACGCCTCCCGTGCTGGACTCCGACGGCTCCTTCTTCCT<br>CTACAGCAAGCTCACCGTGGACAAGAGCAGGTGGCAGCAGGGG<br>AACGTCTTCTCATGCTCCGTGATGCATGAGGCTCTGCACAACCAC<br>TACACGCAGAAGAGCCTCTCCCTGTCTCCGGGTAAATGATTCTAG<br>A |
| LALAPG-3 light chain with signal peptide | GCGGCCGCAAACTACAAGACAGACTTGCAAAAGAAGGCATGCAC<br>AGCTCAGCACTGCTCTGTTGCCTGGTCCTCCTGACTGGGGTGAG<br>GGCCGCTTTGACCCAGCCTGCTTCTGTGTCTGCCAATCTGGGCG<br>GAACCGTGAAGATTACCTGCTCTGGCTCTCGGGGAAGGTACGGT<br>TGGTATCAGCAGAGGAGTCCCGGATCAGCCCCCGTTACTGTGAT<br>TTATAGGGACAATCAGAGACCCTCCAACATTCCCTCTAGGTTCTC<br>CTCCTCCACATCCGGCTCTACATCCACACTGACCATCACCGGCG<br>TGCAGGCTGACGATGAGTCCGTTTACTTCTGCGGCTCCTACGAC<br>GGCTCCATTGACATCTTCGGCGCCGGTACCACCCTCACAGTCTT<br>GCGTACGGTGGCTGCACCATCTGTCTTCATCTTCCCGCCATCTGA<br>TGAGCAGTTGAAATCTGGAACTGCCTCTGTTGTGTGCCTGCTGAA<br>TAACTTCTATCCCAGAGAGGCCAAAGTACAGTGGAAGGTGGATAA<br>CGCCCTCCAATCGGGTAACTCCCAGGAGAGTGTCACAGAGCAGG<br>ACAGCAAGGACAGCACCTACAGCCTCAGCAGCACCCTGACGCTG<br>AGCAAAGCAGACTACGAGAAACACAAAGTCTACGCCTGCGAAGT<br>CACCCATCAGGGCCTGAGTTCGCCCGTCACAAAGAGCTTCAACA<br>GGGGAGAGTGTTGATTCTAGA |
